# A tripartite complex HIV-1 Tat-cyclophilin A-capsid protein enables Tat encapsidation that is required for HIV-1 infectivity

**DOI:** 10.1101/2022.08.07.503104

**Authors:** Malvina Schatz, Laetitia Marty, Camille Ounadjela, Phuoc Bao Viet Tong, Ilaria Cardace, Clément Mettling, Pierre-Emmanuel Milhiet, Luca Costa, Cédric Godefroy, Martine Pugnière, Jean-François Guichou, Jean-Michel Mesnard, Mickaël Blaise, Bruno Beaumelle

## Abstract

HIV-1 Tat is a key viral protein that stimulates several steps of viral gene expression. Tat is especially required for the transcription of viral genes but it is still not clear if and how Tat is incorporated into HIV-1 virions. Cyclophilin A (CypA) is a prolylisomerase that binds to HIV-1 capsid protein (CA) and is thereby encapsidated. Here we found that a Tat-CypA-CA tripartite complex assembles in HIV-1 infected cells. Biochemical and biophysical studies showed that high affinity interactions drive the assembly of this complex. Virions devoid of encapsidated Tat showed a 5-10 fold decrease in HIV-infectivity and, conversely, encapsidating Tat into ΔTat viruses greatly enhanced infectivity. The absence of encapsidated Tat decreases the efficiency of retrotranscription by ∼50% and transcription by 99%. We thus identified a Tat-CypA-CA complex that enables Tat encapsidation and showed that encapsidated Tat is required to initiate robust HIV-1 infection and viral production.

## Introduction

HIV virions contain key viral proteins needed to initiate infection. Indeed, HIV-enzymes such as reverse transcriptase that enables the production of viral cDNA and integrase that will insure insertion of this cDNA into cellular DNA are known to be encapsidated. After viral DNA integration, the next step of the infection cycle is transcription and, surprisingly, HIV transactivating protein Tat that is required for efficient transcription is not thought to be present into HIV virions (Swanson and Malim, 2008). It is thus generally accepted that viral transcription becomes efficient, *i.e*. produces full length viral mRNAs, when neosynthesized Tat becomes available. Indeed, in the absence of Tat, viral transcription is poorly efficient (Feinberg et al., 1991) and it is not clear how Tat can be produced in the absence of Tat (Ott et al., 2011). In addition to transcription, Tat was also found to stimulate retrotranscription (Boudier et al., 2014; Harrich et al., 1997). Biological relevance of this effect can only result from the presence of encapsidated Tat. A proteomic study identified Tat associated with purified HIV virions (Chertova et al., 2006), but the specificity and mechanistic of Tat association to the viral particle remains to be established.

Cyclophilin A (CypA) is a chaperone of the prolylisomerase-immunophilin family. CypA is known to be encapsidated into HIV-1 virions (Franke et al., 1994) due to its affinity (Kd∼16µM) for the G89-P90 motif located within an exposed loop of the capsid protein (CA). The CA protein (24 kDa) is produced during maturation of the Gag precursor polyprotein (55 kDa) by the viral protease. CypA binding to CA results in the incorporation of 200-250 molecules of CypA/virion, *i.e*. ∼1 CypA/ 10 CA. The CA-G89V mutation in CA prevents CypA binding to CA (Yoo et al., 1997). In target cells, CypA binding to CA was found to protect the capsid from restriction by TRIM5α (Kim et al., 2019; Selyutina et al., 2020). Interestingly, viruses deficient in CypA binding were found to be affected in retrotranscription efficiency, but the underlying mechanism remained to be uncovered (De Iaco and Luban, 2014).

Here we show that a tripartite complex Tat-CypA-CA forms in infected cells, that this complex is stable in solution, involves nanomolar affinities and enables efficient Tat encapsidation into HIV virions at the same level as CypA, *i.e.* 200-250 Tat molecules/virion. Encapsidated Tat strongly boosts transcription and also increases the efficiency of retrotranscription. Accordingly, encapsidated Tat stimulates by 5-10 folds HIV-1 infectivity in single round infection assays.

## Results

### Tat-CypA interaction does not require CypA active site

We previously observed using immunoprecipitation that HIV-1 Tat could interact with CypA (Chopard et al., 2018). Here, we first examined whether Tat interaction with CypA relies on CypA catalytic site. To this end, we transfected cells with Tat or HIV-1 Gag as a control, and tested if they could be pulled-down form cell lysates by GST-CypA. We used as a control GST-CypA -H126Q. This point mutation within CypA active site was found to prevent its interaction with its substrates (Foster et al., 2011). Accordingly, while Gag (hence CA) could be efficiently pulled down from cell lysates by GST-CypA, it did not interact with GST-CypA-H126Q (Fig.1A). Surprisingly, the efficiency of Tat interaction with CypA-H126Q was ∼50 % of the efficiency of Tat binding to WT CypA, while Tat did not interact with GST. These data indicated that Tat can bind CypA independently of CypA active site, and suggest that a Tat-CypA-CA complex can exist within infected cells.

**Figure 1.**
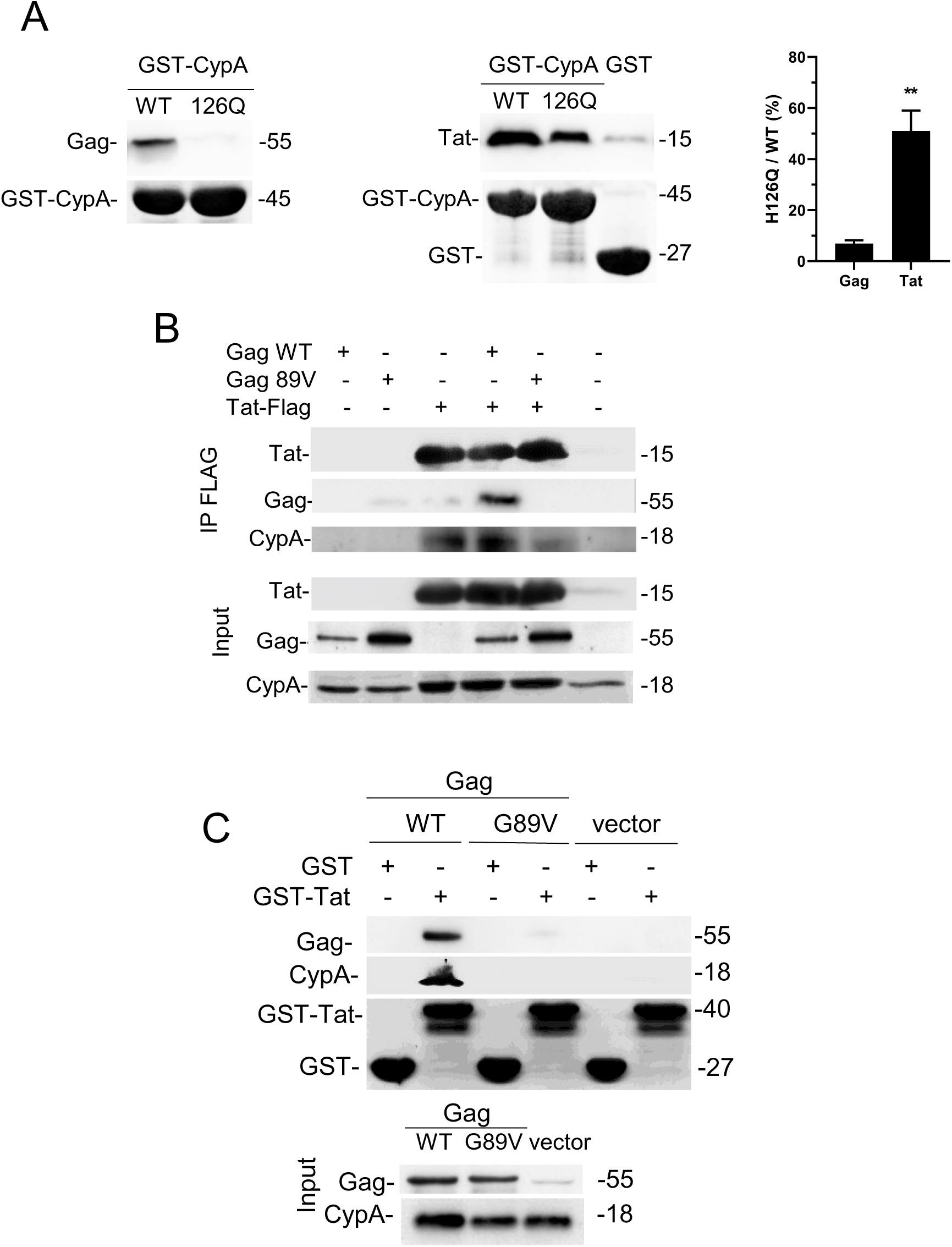
Tat interaction with CypA does not require CypA active site and allows the formation of a Tat-CypA-CA tripartite complex. **A,** HEK 293T cells were transiently transfected with a Gag (left panel) or Tat (right panel) vector. After 36 h cell lysates were incubated with GST, GST-CypA or GST-CypA-H126Q immobilized on GSH-agarose, before washes and anti-Tat, -p24 or-GST Western blots. GST blots showed equal GST loading. The graph shows the quantification of the amount of Tat or Gag pulled down by H126Q/WT GST-CypA (%). Data are mean ± SEM from n=5 experiments. **, p< 0.01 (Student’s t-test). **B**, Immunoprecipitation of the Tat-CypA-CA complex. Cells were transfected with Tat-FLAG, Gag WT, Gag-CA-G89V or empty vector as indicated. After 36 h, Tat-FLAG was immunoprecipitated from cell lysates using an anti-FLAG antibody, before Western blots. **C**, GST pull down by GST-Tat. Cells were transfected with Gag WT, Gag-CA-G89V or empty vector as indicated. After 36 h cell lysates were incubated with GST-Tat immobilized on GSH-agarose, before washes and anti-p24 or-GST Western blots. GST staining showed similar GST loading. **Figure supplement-source data.** Raw immunoblots and datasheet for the graph.

### A tripartite Tat-CypA-CA complex exists within cells

We then examined whether a tripartite Tat-CypA-CA could be observed within transfected cells. We first used Tat-FLAG and anti-FLAG immunoprecipitation to isolate this complex from Tat-FLAG and Gag transfected cells. As shown earlier, CypA is co-immunoprecipitated with Tat (Chopard *et al*., 2018), whether Gag is present or not (Fig.1B). When Tat-FLAG and Gag were cotransfected, WT Gag but not Gag-CA-G89V was immunoprecipitated with Tat. These results suggested that a Tat-CypA-CA complex exists within cells and that, in this complex, CA and CypA interact *via* the well characterized binding of CypA active site to CA-Pro90 (Yoo *et al*., 1997).

To confirm these data, we tried to precipitate the complex using GST-Tat and extracts from Gag transfected cells. GST-Tat, but not GST alone enabled to pull-down CypA and WT Gag (Fig.1C). Gag-CA-G89V could not be pulled-down by GST-Tat.

Altogether, these pull-down experiments strongly suggested that a Tat-CypA-CA complex is formed intracellularly.

### Tat-CypA-CA complex can be purified by gel filtration

We then assessed whether this complex could form using purified recombinant proteins, if it was stable in solution and purifiable. We first analyzed the different combination of the three highly purified proteins by size exclusion chromatography (SEC). CA and CypA alone behaved well and eluted from the SEC column as sharp peaks at ∼14 ml and ∼17 ml (Fig.2A), while Tat eluted as a very shallow peak at 19-20 ml (arrow in Fig.2A top panel). When CA and CypA were mixed, the CA elution peak shifted to earlier retention times, indicating that a CA-CypA complex slightly but significantly bigger than CA alone was formed (Fig.2A and B). This shift was not observed when CypA-H126Q was used indicating that this complex is a *bona fide* CA-CypA complex in which CypA is bound to CA via its catalytic site. When Tat, CA and CypA were mixed then injected, the CA peak did not appear earlier compared to when CA+CypA were analyzed, but when this fraction was analyzed by western blots we found that Tat was recruited at the level of this peak (Fig.2C). CypA-H126Q did not allow the recruitment of Tat to the CA peak. This SEC analysis thus confirmed the existence of the Tat-CypA-CA tripartite complex and indicated that it is stable, can be purified and amenable to biochemical characterization.

**Figure 2.**
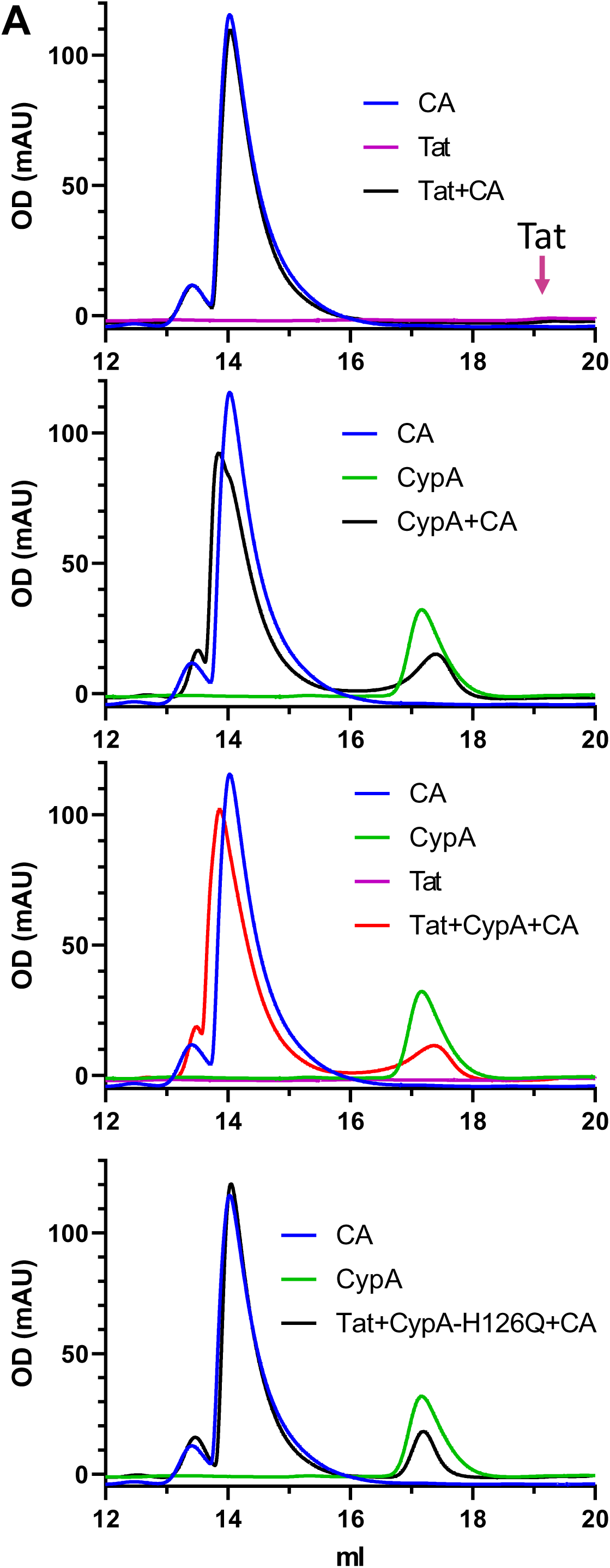
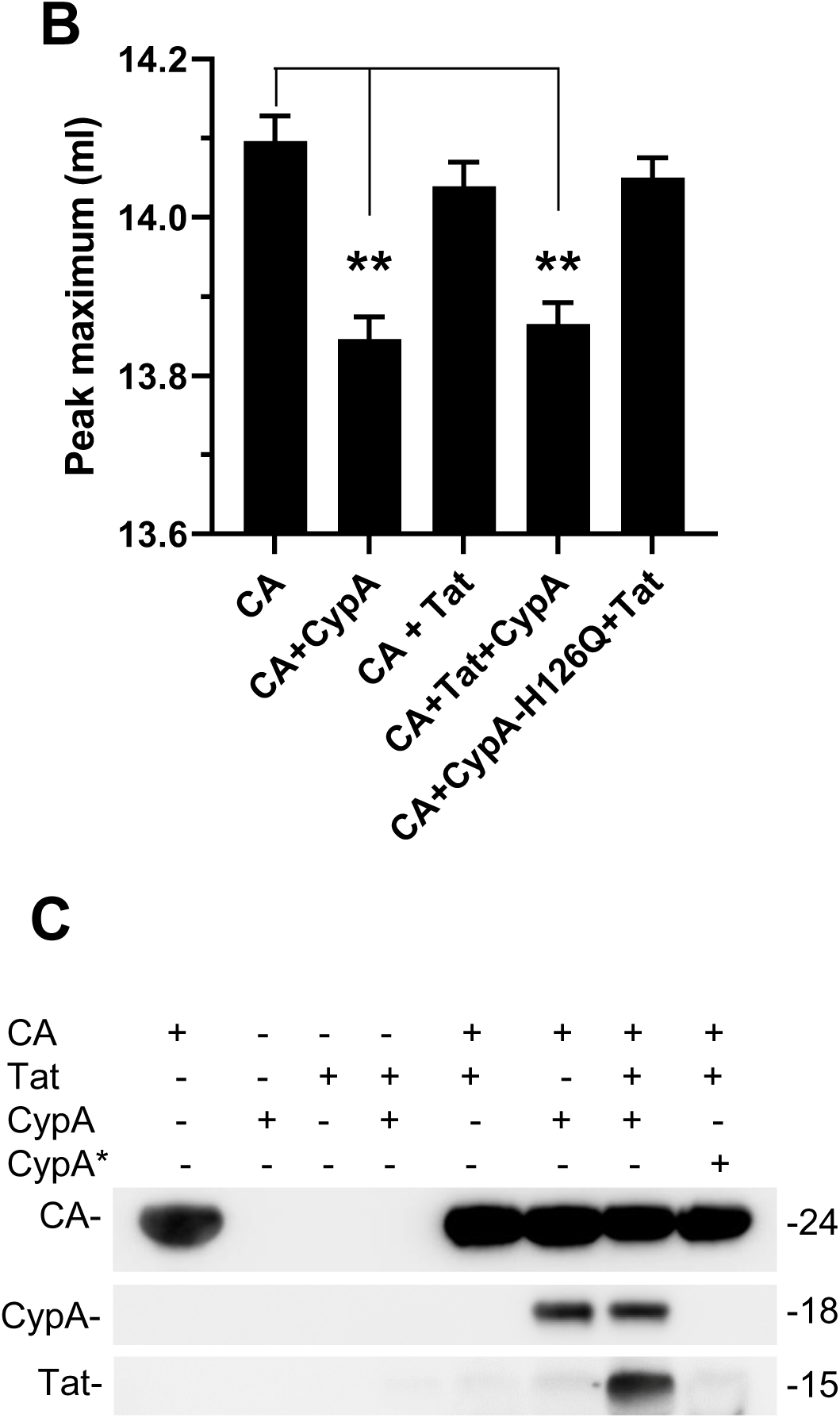
Analysis of the CA-CypA-Tat complex by size exclusion chromatography. The indicated protein mixture (CA, CypA, Tat or the indicated combination) was incubated for 30 min at RT then loaded on a Superdex 200 Increase column. **A**, protein elution profile monitored using OD_280_. **B**, The elution volume corresponding to the maximum OD_280_ of the first peak (CA peak∼ 14 ml) is shown. Means ± SEM of n= 2-3 injections. **, p<0.01 (One Way ANOVA). **C**, fractions corresponding to the capsid peak (∼14 ml) were analyzed by western blot. CypA* is CypA-H126Q. **Figure supplement-source data.** Raw immunoblots and datasheets for graphs.

### Analysis of the Tat-CypA-CA complex by microscale thermophoresis

To further characterize the complex and determine its interaction strengths we determined the dissociation constant at the equilibrium (*K*_d_) of the three partners involved in the Tat-CypA-CA complex by microscale thermophoresis (MST). CypA, the central protein of the complex was fluorescently labeled. Labeled CypA interacted with CA with a K_d_ of 8.3 ± 1.7 µM (Fig.3A) similar to the one reported before for the CypA-CA interaction (Yoo *et al*., 1997), therefore indicating the functionality of fluorescent CypA. A preincubation of CypA with 1 µM Tat did not significantly affect CA-CypA interaction (K_d_ of 5.5 ±0.25 µM). Tat affinity for CypA was higher (K_d_ of 243 ± 23 nM) than that of CA for CypA. This is consistent with the observation that Tat-CypA interaction involves CypA regions that are outside the active site (Fig.1). Indeed, as a generalist chaperone, CypA exhibits a poor substrate specificity. The best substrates are those such as HIV-1 CA that contains the Gly-Pro motif (Howard et al., 2003) and their K_d_ is > 5 µM (Piotukh et al., 2005). Tat submicromolar affinity for CypA thus confirms that the ligand is not only interacting with CypA active site. Interestingly, Tat affinity for CypA was enhanced 5-fold (K_d_ of 55 ± 14 nM) when CypA was preincubated with 20 µM CA.

**Figure 3.**
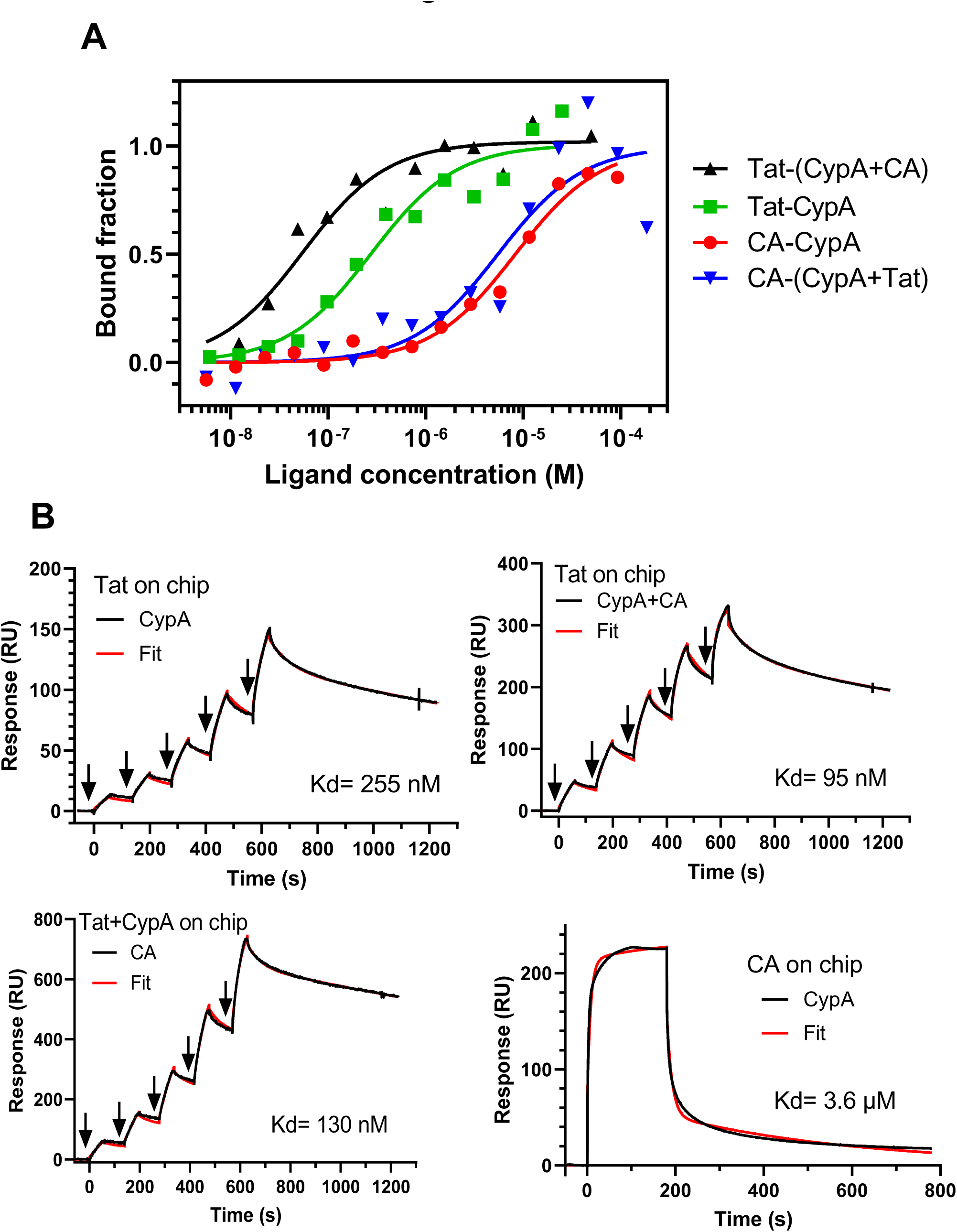
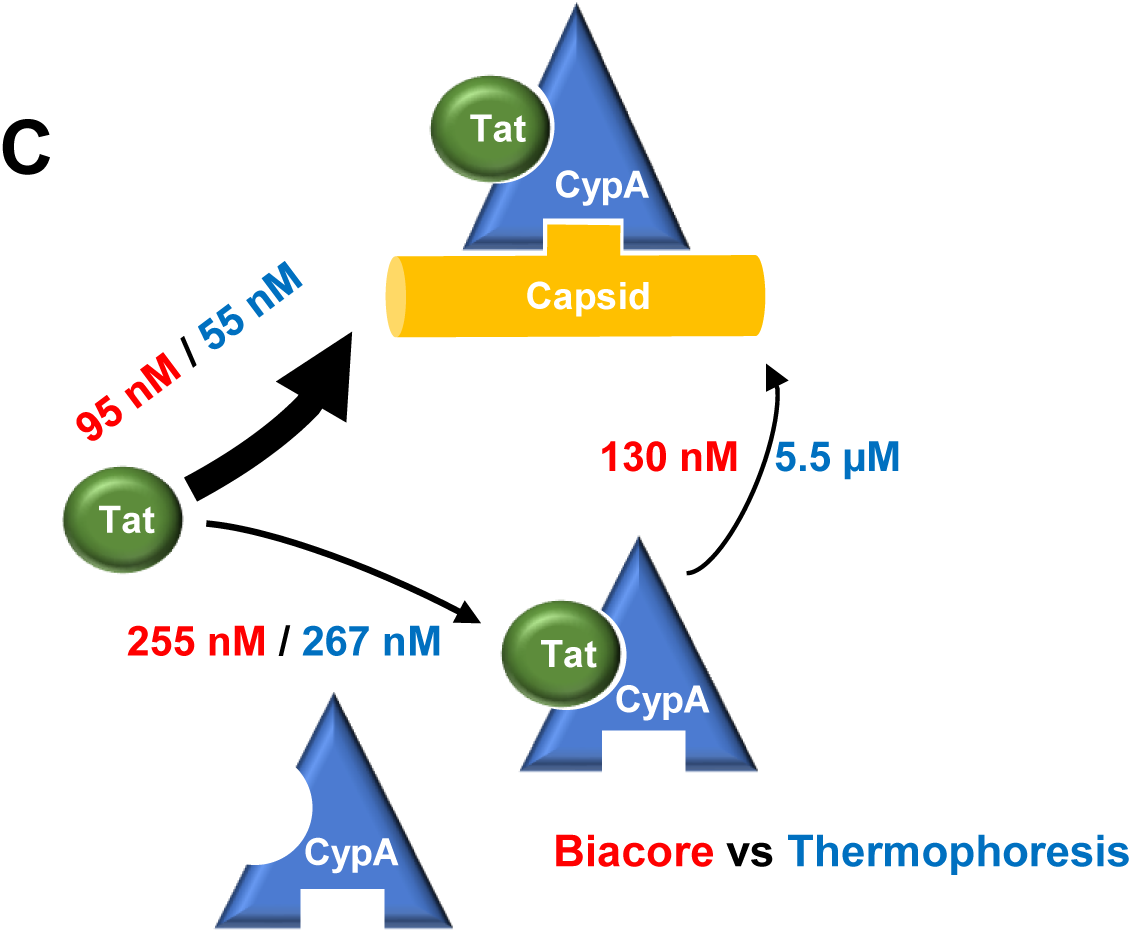
Analysis of the interactions involved in the assembly of the Tat-CypA-CA complex. **A**, analysis by MST. CypA was fluorescently labeled and its interaction with increasing concentrations of Tat or CA was tested. When indicated by (CypA+CA) or (CypA+Tat) fluorescent CypA (50 nM) was first mixed with a saturating concentration of CA (20 µM) or Tat (1 µM) before adding escalating concentrations of the third partner. **B,** analysis by SPR. Tat was immobilized on the Biacore chip before applying the indicating partner using escalating concentrations. CypA or premixed CypA+CA were applied at 0.37 / 0.75/ 1.5 / 3 / 6 µM. When indicated (Tat+CypA on chip), CypA (4 µM) was applied to immobilized Tat before applying CA (0.37 / 0.75/ 1.5 / 3 / 6 µM) to monitor CA binding to CypA presented by Tat. Arrows indicate injections. **C**, Graphical abstract of the affinities involved in the assembly of the Tat-CypA-CA complex as observed using MST and SPR. **Figure supplement-source data.** Datasheets for graphs.

### Analysis of the Tat-CypA-CA complex using surface plasmon resonance

To confirm MST data we studied assembly of the Tat-CypA-CA complex using surface plasmon resonance (SPR). When Tat was immobilized on the chip before applying increasing doses of CypA, very strong interaction in the nM order was observed (K_d_ of 255 ± 10 nM; Fig.3B), in very good agreement with MST data. We did not observe any binding of CA to Tat (Figure S1) while when a CypA+CA complex was applied to a Tat chip a K_d_ of 95 nM was observed (Fig.3B) indicating, as observed using MST, that Tat binds with more affinity to the CypA-CA complex than to CypA alone. The CA-CypA interaction has been studied extensively and, in agreement with previous measurements, using SPR(Yoo *et al*., 1997) we observed a K_d_ of ∼3.6 µM for CypA binding to CA. Nevertheless, when we followed CA binding to the Tat-CypA complex assembled on the chip, a K_d_ of 110 ± 20 nM was obtained. Hence, SPR data indicated that the presence of Tat on CypA facilitates the CA-CypA interaction, while this was not observed using MST. Both techniques otherwise provided very consistent affinity constants (Fig.3C). They both indicated that Tat is preferentially recruited by CypA when CypA is already bound to CA. This suggests that Tat is preferentially recruited where CypA is bound to CA, *i.e.* at HIV budding sites, and therefore that the Tat-CypA-CA complex can drive Tat encapsidation.

### Tat is encapsidated in a CypA-dependent manner

To examine whether the Tat-CypA-CA complex enables Tat encapsidation, we first purified HIV virions from transfected HEK 293T cells. Tat WT was observed in this purified virus preparation using two different monoclonal antibodies, a first one that recognize the N-ter (res 1-9) of the protein and a second that binds to a conformational epitope (Mediouni et al., 2012). A Tat mutant (Tat-W11Y) that poorly binds PI(4,5)P_2_ (Rayne et al., 2010) was also associated with virions, it was just not detected by the anti-Tat that recognizes the N-ter of the protein probably because the W11Y mutation severely inhibits the binding of this antibody (Fig.4A). Hence, although HIV-1 budding takes place from PI(4,5)P_2_ enriched area and virions are accordingly enriched in this phosphoinositide (Sundquist and Krausslich, 2012), these results indicates that Tat binding to PI(4,5)P_2_ is not required for Tat association with purified virions. Non palmitoylable Tat (Tat-C31S (Chopard *et al*, 2018)) was also encapsidated, although this mutant was also less efficiently recognized by the N-ter anti-Tat. This result indicated that Tat palmitoylation is not required for Tat encapsidation. Very low amounts of Tat could be detected in CA-G89V virions even upon overexposure and the amount of Tat associated with these virions usually represented less than 5% of that present in WT virions. In agreement with previous studies (Schaller et al., 2011; Sokolskaja et al., 2004), viruses with CA-G89V did not incorporate CypA. The immunophilin FKBP12 that is weakly encapsidated (∼25 copies/virions (Briggs et al., 1999)) and that can interact with Tat (Chopard *et al*., 2018) was also present in CA-G89V virions. Altogether these results indicated that, in agreement with biochemical data, Tat encapsidation relies on the presence of encapsidated CypA, while Tat interaction with PI(4,5)P2 or FKBP12 does not seem to allow significant Tat encapsidation. This CypA-dependent Tat encapsidation also indicates that the presence of Tat in purified virus preparation is not the result of contamination by exosomes or other cell-derived vesicles. Tat was quantified using semiquantitative blots (Figure S2). A calculation based on p24 ELISA of purified virions and the number of 2000-2500 CA per virion (Freed, 2015) showed that 200-250 Tat molecules were present by virion, just as it is the case for CypA (Yoo *et al*., 1997). This result is consistent with the high affinity of Tat for CypA-CA (K_d_ 55-95 nM; Fig3C) compared to the 38-150 lower affinity of CypA for CA (3.6- 8.3 µM; Fig3A-B). Hence, the limiting factor for Tat encapsidation is CypA binding to CA, and not Tat binding to CypA.

**Figure 4.**
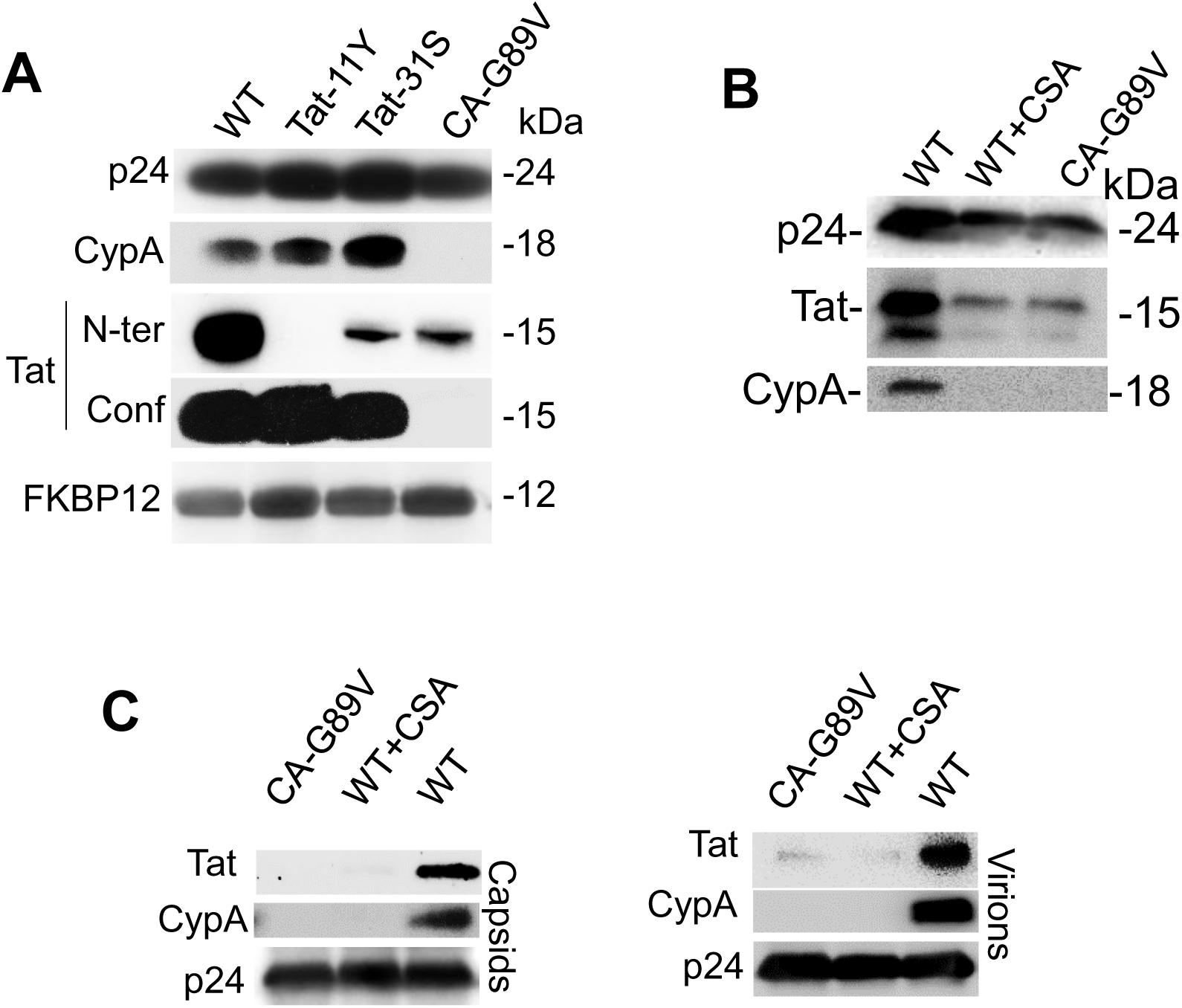
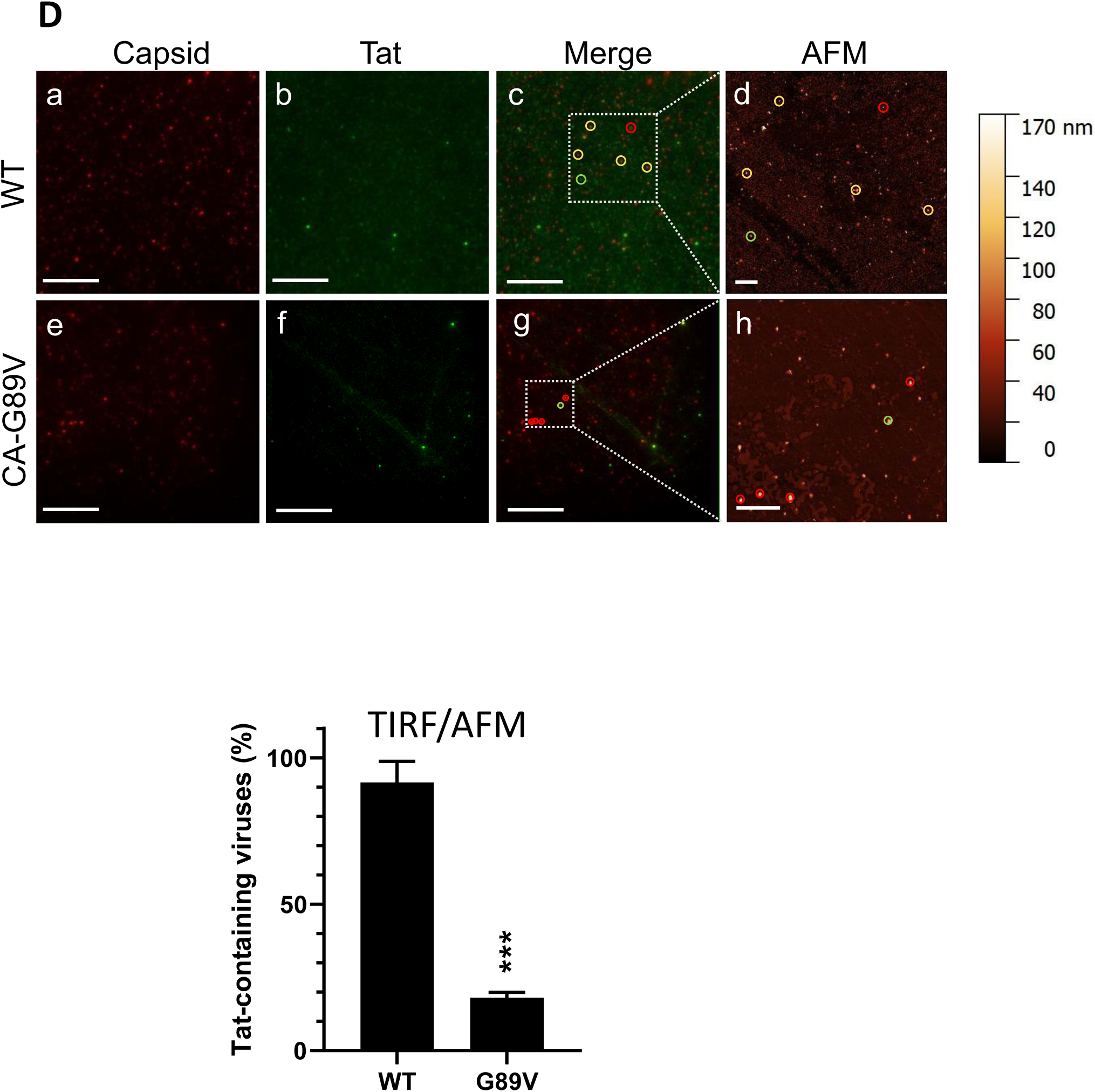
HIV-1 Tat is encapsidated in a CypA-dependent manner. **A**, Transfected cells. HEK cells were transfected with pNL4.3, either WT, Tat-W11Y, Tat-C31S or CA-G89V as indicated. After 48 h, viruses were purified from the supernatant by ultracentrifugation through a sucrose cushion, before analysis by Western blots. Two different antibodies were used for Tat, one that recognizes the N-ter of the protein (res1-9) and a second whose epitope is conformational (Conf). Tat-W11Y was not recognized by the antibody against the N-ter of the protein. Overexposed gels only enabled Tat detection in CA-G89V virions, that are devoid of CypA. **B,** Infected cells. HIV_NL4.3_ WT or CA-G89V was used to infect Jurkat-CD4-CCR5 cells and infection was allowed to proceed for 9 days adding fresh cells every 3 days before harvesting virions for analysis. When indicated 2 µM CSA was added to cells to prevent CypA encapsidation. **C,** Capsids from WT, CSA or CA-G89V viruses were purified following Triton X-100 treatment to solubilize the viral envelope and both purified viruses and capsids were analyzed by Western blots, **D,** purified WT or CA-G89V virions were coated on coverslips and labelled for p24 (red) and Tat (green) before AFM and TIRF imaging. Images were aligned using Ec-CLEM. Viruses were identified as AFM spots >100 nm. AFM-identified virions are circled in red, green or yellow if they are positive for CA, Tat or both, respectively. Scale bars are 10 µm and 2 µm for fluorescence and AFM images, respectively. Representative images are shown. The graph corresponds to the percentage of viruses AFM^+^ and CA^+^ containing Tat for WT and CA-G89V virions (n=3 from entirely independent experiments. mean ± SEM; Student’s t-test ***, p<0.001). **Figure supplement-source data.** Datasheets for graphs, summary statistics and raw immunoblots.

It was important to validate this CypA-dependent Tat encapsidation observed using transfected HEK 293T cells using viruses obtained following infection. We thus analyzed HIV-1 virions purified from infected Jurkat T-cells (Fig.4B). For these viruses produced by infected T-cells, Tat was present in WT viruses as a doublet. While the major band represents full-length Tat (two-exon) of 101 residues, the minor band probably corresponds to the single-exon form of Tat (72 residues (Verhoef et al., 1997)) that is also produced during infection (Karn and Stoltzfus, 2012). Tat incorporation was strongly decreased when CypA encapsidation was inhibited by the CA-G89V mutation or when infected cells were treated with cyclosporin-A (CSA) a CypA inhibitor that prevents its interaction with CA(Luban et al., 1993) and CypA encapsidation (Franke *et al*., 1994; Thali et al., 1994). The result that both two- and single-exon Tat can be encapsidated suggests that Tat encapsidation motif, *i.e.* Tat CypA-binding site, resides within the first 72 residues of the protein.

To further check whether Tat was associated with HIV capsids we purified capsids form purified HIV-1 virions treated with TX-100 to dissolve the viral envelope (Xu et al., 2020). Tat association with purified HIV-1 capsids was observed for WT viruses but only weakly for CA-G89V or viruses from CSA-treated cells that are devoid of CypA (Fig.4C).

Altogether, these biochemical data indicated that Tat is encapsidated by HIV-1 in a CypA-dependent manner.

To confirm these biochemical observations, we used a morphological approach. We examined virions using a correlative atomic force / total internal reflection fluorescence microscopy (AFM/TIRF) setup ((Dahmane S et al., 2019). To this end, freshly purified virions plated on coverslips were permeabilized for immunofluorescence labeling of Tat and CA. HIV-1 virions have a diameter of 100-120 nm (Freed, 2015). Virions were identified by AFM as round-shaped particles with height > 100 nm and fluorescent CA labeling. Tat was observed in >90% of WT virions but only of 18% of CA-G89V viruses (Fig.4D).

Altogether, biochemical and morphological data showed that Tat is encapsidated in CypA-dependent manner.

### Encapsidated HIV-1 Tat is delivered to the cytosol

We then assessed whether encapsidated Tat is delivered to the cytosol upon infection. To examine this point, we transfected human primary CD4 T-cells with an LTR-firefly luciferase vector whose transcription will be activated by Tat and a renilla control vector, before adding VSV-G pseudotyped ΔEnv viruses. To specifically follow the cytosolic delivery of encapsidated Tat and prevent Tat synthesis by infected cells, AZT was used to inhibit reverse transcription. Transactivation by Tat was followed using the firefly/renilla activity ratio (Fig.5A).

**Figure 5.**
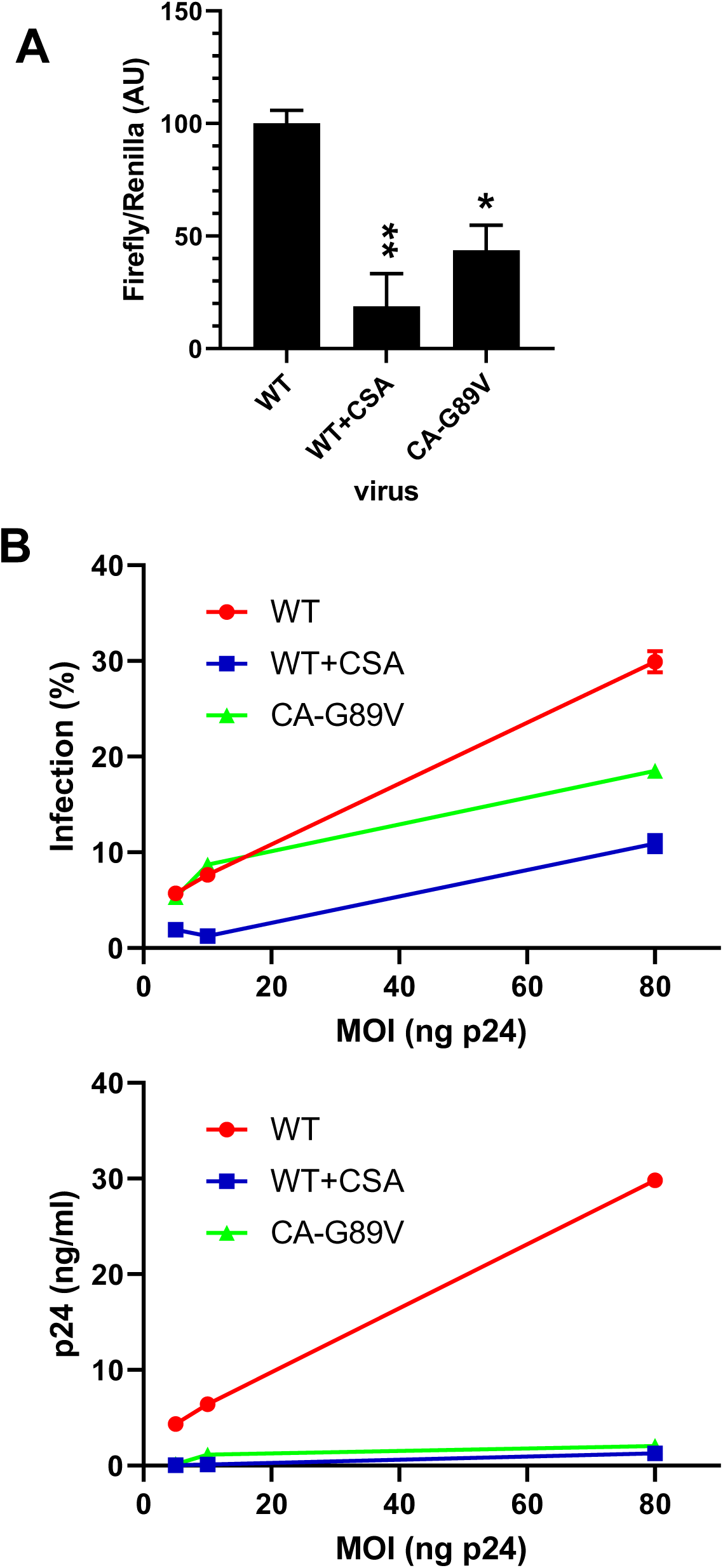
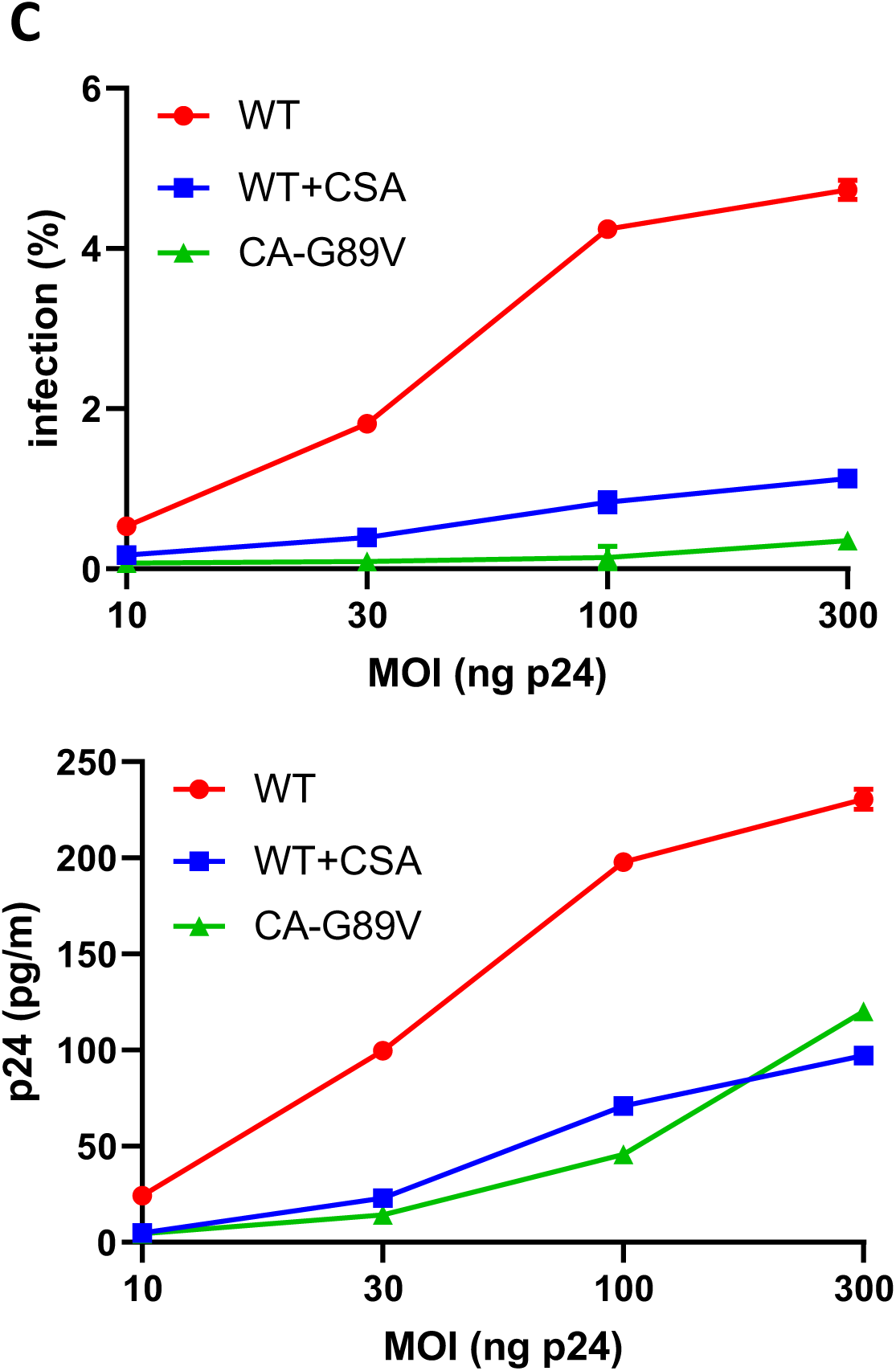
Encapsidated HIV-1 Tat is delivered to the cytosol and is required for efficient HIV infection. **A**, Cytosolic delivery assay. Human CD4 primary T-cells were transfected with LTR-firefly luciferase and TK-renilla luciferase (control vector). After 18 h, cells were infected with the pseudotyped HIV_NL4.3_, either WT or CA-G89V or prepared in the presence of 2 µM CSA. Infections were performed in the presence of 25 µM AZT to inhibit Tat neosynthesis. Cells were lysed 18 h pi to assay luciferase activities. The graph shows the Firefly/Renilla activity ratio (mean ± SEM; n=3). One Way ANOVA results *, p<0.05; **, p<0.01. **B** and **C,** single-round infections. Jurkat T-cells (**B**) and human primary CD4 T-cells (**C**) were infected with the indicated amount (ng p24) of VSV-G pseudotyped HIV_NL4.3_ viruses. No CSA was added, it was used during viral production by HEK cells only. After 36h, cells were stained for p24 before FACS analysis to determine the efficiency of infection (%), and virus concentration in the cell medium was assayed using p24 ELISA (means ± SEM; n=2). Representative experiments from n=3. Most error bars are within the symbol size. **Figure supplement-source data.** Datasheets for graphs and summary statistics.

The absence of encapsidated Tat induced a ∼60% or ∼80 % decrease in firefly production for CA-G89V viruses or viruses from CSA treated cells, respectively. This result indicated that Tat encapsidated by WT viruses is delivered to the cytosol upon infection and is transcriptionally active.

### Encapsidated Tat is required for HIV-1 infectivity

Since encapsidated Tat is delivered to the cytosol and transcriptionally active, it should enable the virus to establish infection more quickly and efficiently. We examined the effect of encapsidated Tat on infection by HIV-1 using single-round infectivity assays (McMahon et al., 2009). To this end, Jurkat T-cells were infected using VSV-G pseudotyped ΔEnv viruses. Results from p24 FACS analysis of cells showed that for the highest MOI, cell infection was reduced by 60% or 40% when Tat was removed from virions using CSA or CA-G89V, respectively (Fig.5B). When viral production was monitored using p24 ELISA a reduction of 95% and 80% for CSA or CA-G89V viruses was observed (Fig.5B). When human primary CD4 T-cells were used as targets stronger differences were observed since whatever the MOI, viruses without encapsidated Tat were ∼five-fold (CSA) or more than 10-fold (CA-G89V) less infectious than WT HIV-1 (Fig.5C). The reduced toxicity of viruses with the CA-G89V or prepared from CSA treated cells was reported earlier by different groups (Sokolskaja *et al*., 2004; Towers et al., 2003). The absence of CypA on CA-G89V capsids triggers restriction by TRIM5α, but this effect is only observed in primary T-cells (Kim *et al*., 2019) and not in transformed cells (Sokolskaja et al., 2006) such as Jurkat cells (Selyutina *et al*., 2020). Hence, it is likely that the CA-G89V mutant infects less efficiently primary CD4 T-cells than the Jurkat cell line due to restriction by TRIM5α.

Altogether single-round infection assays thus indicated that encapsidated Tat is required to establish both efficient infection and sustained viral production.

### Encapsidated Tat facilitates viral reverse transcription

We then examined at which stage encapsidated Tat could act to favor infection. Regarding early steps of the infection cycle, Tat was found to facilitate reverse transcription (Boudier *et al*., 2014; Harrich *et al*., 1997; Lalonde et al., 2011) and CypA was found to be required for the initiation of reverse transcription (Braaten et al., 1996). We therefore examined whether encapsidated Tat affected the production of early and late reverse transcription products. We quantified these products by qPCR at early time points following infection (6 and 10h), when their production is maximum (Butler et al., 2001). The absence of encapsidated Tat in virions significantly decreased the efficiency of reverse transcription by ∼50% (Fig.6A).

**Figure 6.**
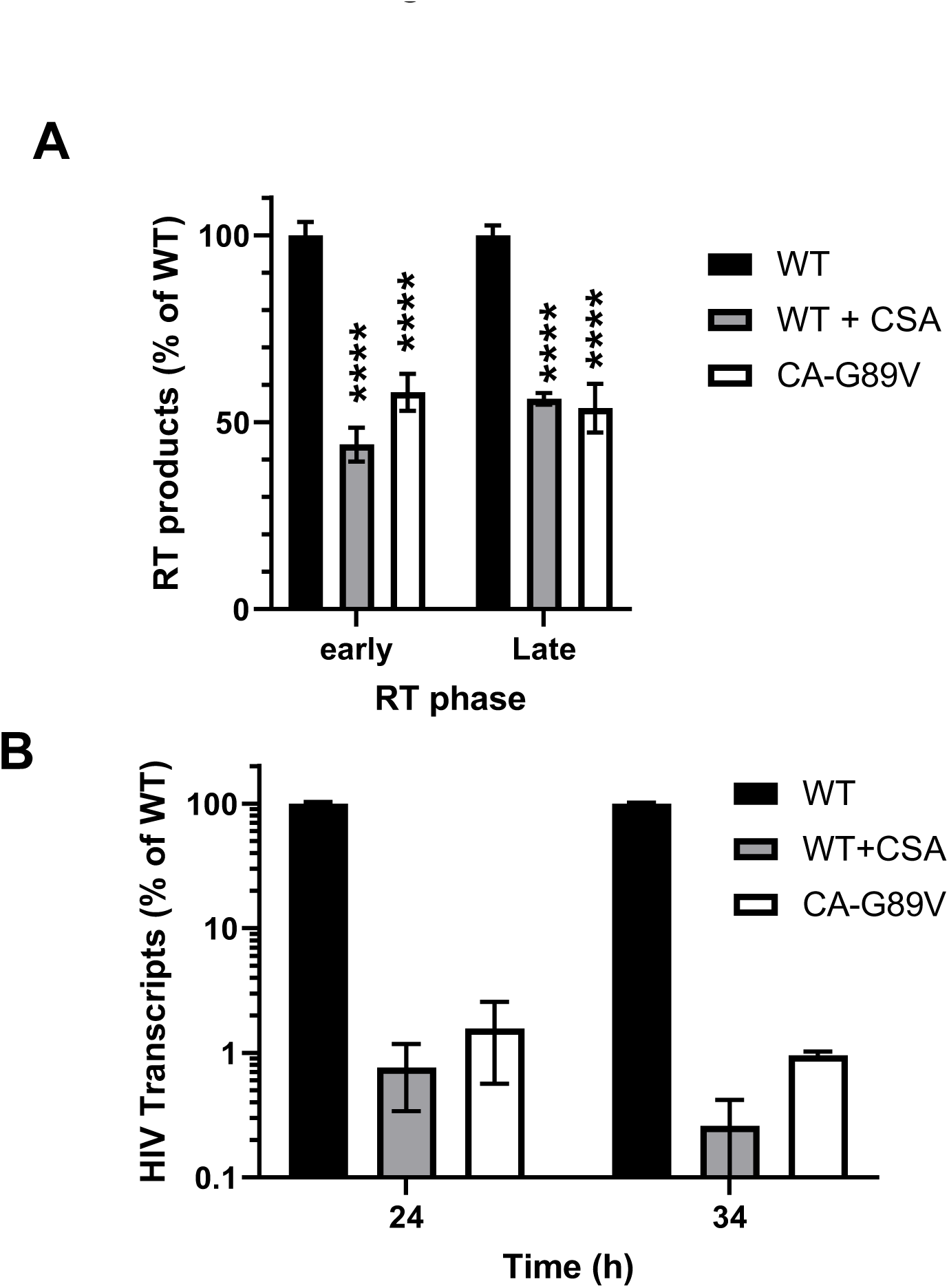
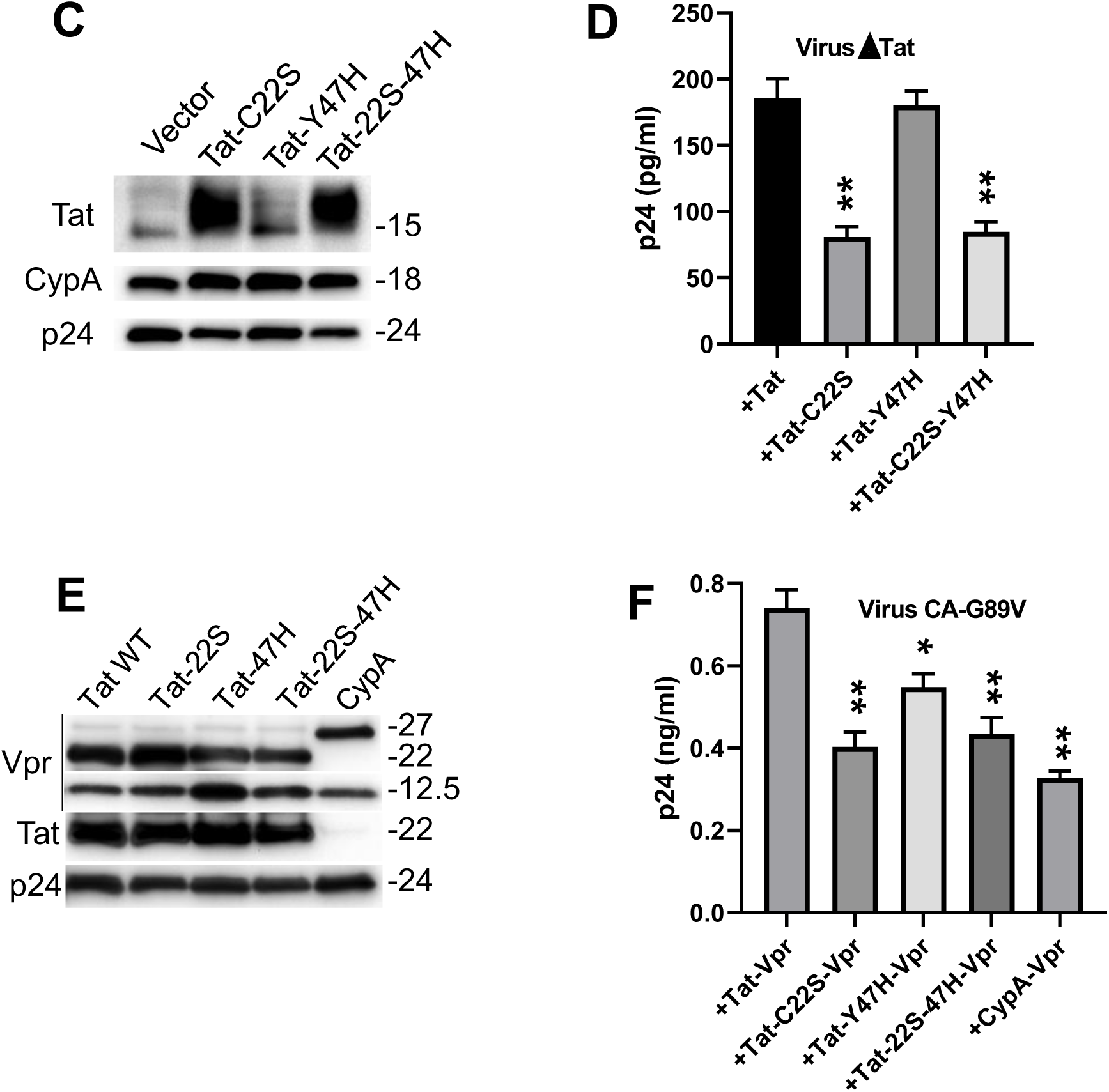
Encapsidated Tat increases reverse transcription and strongly stimulates transcription. **A**, Quantification of early and late RT products. Jurkat T-cells were infected with HIV_NL4.3_, either WT or CA-G89V or prepared in the presence of 2 µM CSA. Cells were harvested 6h after infection for DNA extraction and qPCR. **B,** q-RT-PCR of viral RNA. Cell RNA was extracted 24h or 34h after infection. It was reversed transcribed to cDNA for qPCR. All data are presented as % of WT virus data. PCR data are means ± SEM of 2 independent experiments each with n=4. **C**, ΔTat ΔEnv pNL4.3 was cotransfected in HEK293T cells with VSV-G to pseudotype virions and a vector coding for Tat WT, C22S, Y47H or 22S-47H as indicated. Tat mutants were cotransfected with WT Tat (ratio 1 WT/ 2 mutant) to insure viral production by Tat-22S. Viruses were purified for biochemical analysis by Western blot. **D,** Tat-complemented viruses were used to infect Jurkat cells in single-round infection assays and viral production in the medium was monitored using p24 ELISA. **E,** CA-G89V pNL4.3 was cotransfected in HEK293T cells with VSV-G to pseudotype and a vector coding for the indicated Vpr chimera. Viruses were purified for biochemical analysis by Western blots. Anti-Vpr detects the endogenous viral Vpr at 12.5 kDa, Tat-Vpr at 22 kDa and CypA-Vpr at 27 kDa. **F,** CA-G89V viruses with Vpr chimera were used to infect Jurkat cells in single-round infection assays and viral production in the medium was monitored using p24 ELISA. Data are means ± SEM (n=2) of a typical experiment that was repeated twice. One Way ANOVA *, p<0.05; **, p<0.01. **Figure supplement-source data.** Datasheets for graphs, summary statistics and raw immunoblots.

### Encapsidated Tat strongly stimulates transcription

Tat is required for the efficient transcription of viral genes (Ott *et al*., 2011). We examined by qRT-PCR of intracellular viral RNA to what extend encapsidated Tat could affect viral transcription at 24 h and 34 h post-infection, *i.e.* following the integration of viral DNA (Butler *et al*., 2001). Viruses without encapsidated Tat (CSA or CA-G89V) were strongly affected in viral transcription whose efficiency was ∼1% of that of WT viruses (Fig.6B).

### Complementation of Tat-deficient viruses confirms the role of encapsidated Tat in infection

The CSA and CA-G89V viruses are devoid of encapsidated Tat but also of CypA and, furthermore, the CA-G89V mutant is unable to bind CypA. This mutation of the CypA binding loop was reported to affect several early steps of HIV-1 infection cycle, such as recognition by TRIM5α (Kim *et al*., 2019; Selyutina *et al*., 2020), but also nuclear import (Lahaye et al., 2016). To specifically examine the role of encapsidated Tat in infection, we prepared viruses with CypA and different versions of Tat either WT, or with a point mutation reported to selectively inactivate transcription (Tat-C22S (Jeang et al., 1999)) or reverse transcription (Tat-Y47H (Apolloni et al., 2003)). According to an RMN-based structure (Bayer et al., 1995) the mutated residues are, except for the hydroxyl group of Y47, located inside the protein. Their mutation was thus unlikely to affect CypA binding and Tat encapsidation. Viruses were produced using a ΔEnv ΔTat viral clone that was cotransfected in HEK 293T cells with Tat and VSV-G vectors. The Tat-C22S and Tat-47H WT mutants were cotranfected 2/1 with WT Tat to insure significant viral production by Tat-C22S virions.

Biochemical analyses of the virions showed that they contained equivalent amounts of CA and CypA. Tat-Y47H was encapsidated as efficiently as WT Tat, while Tat-C22S, that exhibits a change in electrophoretic mobility, was more efficiently encapsidated (Fig.6C). We monitored viral production by these pseudotyped viruses using single round infection assays. While the Tat-C22S mutation decreased viral production by 50-60%, the Tat-Y47H mutation did not significantly affect production (Fig.6D). Virions with the double mutant Tat-22S-47H behave as those with Tat-C22S. These observations are consistent with the data indicating that Tat encapsidation strongly stimulates transcription and only marginally increases reverse transcription.

To confirm this loss of function data, we used a gain of function approach. To this end, we prepared CA-G89V viruses in which we encapsidated Tat using Vpr fusions. Fusion with the viral protein Vpr that is efficiently encapsidated is an efficient method to encapsidate proteins into HIV virions (Yao et al., 1999). We prepared Tat-Vpr fusions using WT, C22S, Y47H or 22S-47H versions of Tat, and CypA-Vpr was used as a negative control. Biochemical analysis of the virions showed that Vpr chimera were encapsidated to similar levels (Fig.6E). In single-round infections, the highest viral production was observed for the virions with Tat-Vpr (Fig.7D). The negative control CypA-Vpr level represents the participation of neosynthesized Tat in viral production. Viral production due to Vpr-Tat was essentially ablated by the Tat-C22S mutation, while the Tat-47H mutation had a milder effect on viral production. The double mutant Tat-22S-47H behaved essentially as the Tat-C22S transcription inactive mutant.

Altogether, results from qRT-PCR data obtained with CSA or G89V viruses, together with single-round infection data of ΔTat complemented viruses and VPR-chimeras confirmed that encapsidated Tat enables to efficiently boost transcriptional activity. Encapsidated Tat also acts earlier by increasing the reverse transcription activity but the three approaches indicated that this effect is weaker compared with the strong increase in Tat-catalyzed transcription.

## Discussion

The CypA-binding loop within HIV capsid protein has several key roles within the infected cell. First, CypA binding protects HIV-1 core from the TRIM5α restriction factor (Kim *et al*., 2019; Selyutina *et al*., 2020). The presence of CypA on the incoming capsid is also essential for the action of the nuclear envelope protein SUN2 that is needed for HIV infection (Lahaye *et al*., 2016). The CypA binding loop from CA is also involved in capsid nuclear import by binding to the cytoplasmic nuclear pore complex protein Nup358/RanBP2 (Schaller *et al*., 2011) and transportin-1 (Fernandez et al., 2019). Here we document another role of this CypA-binding loop, that acts at the level of nascent virions by enabling Tat encapsidation *via* the formation of a CA-CypA-Tat tripartite complex. Some reports indicated that Tat could be associated with HIV virions. Tat was detected in a proteomic study of purified HIV viruses, but no quantification was made (Chertova *et al*., 2006). A potential caveat with Tat is that it is actively secreted by infected cells (Ensoli et al., 1990; Rayne *et al*., 2010) and that Tat association with viral particles could therefore take place after viral budding and be potentially non-specific. Nevertheless, a gp120-Tat high affinity interaction was observed (Marchio et al., 2005). The presence of Tat specifically or unspecifically bound to virion surface might be responsible for the presence of Tat at low levels in viruses from CA-G89V mutants or CSA-treated cells, but this is a very minor fraction of virion-associated Tat that is encapsidated in a CyA-dependent manner (Fig.4).

Throughout this study we monitored HIV infectivity using single-round assays to specifically examine the role of encapsidated Tat in the initial steps of infection. Although we could confirm the role of Tat (Boudier *et al*., 2014; Harrich *et al*., 1997; Lalonde *et al*., 2011) and CypA (Braaten *et al*., 1996) on reverse transcription, the effect of encapsidated Tat on the RT step was less pronounced than that on the transcription of viral genes that enables to efficiently initiate viral production before neosynthesized Tat becomes available. We observed that 200-250 copies of Tat are associated with virions. This number of molecules in the human T-cell volume of 176 µm^3^ (Chapman et al., 1981) generates a Tat concentration of ∼2 nM, above the Kd (0.4-0.8 nM) involved in the assembly on HIV transcription complex Tat-pTEFb-TAR (Zhang et al., 2000). Each HIV virus therefore encapsidates enough Tat to efficiently stimulate transcription. While recent data showed that HIV-1 capsid remains intact until it enters the nucleus (Zila et al., 2021), live-cell imaging indicated that CypA dissociates from the capsid upon nuclear entry (Francis and Melikyan, 2018). Hence, upon HIV entry, Tat should be delivered within or at the nuclear doorstep so that Tat is present close to its action site.

Here we show that, by encapsidating Tat, HIV can take control of transcription from the beginning, so that this key step of viral gene expression is efficient as early as possible. The virus thus encapsidates the three viral proteins, reverse transcriptase, integrase and Tat that are needed to initiate the production of viral RNA and proteins.

## Materials and methods

### Materials

Opti-MEM, Lipofectamine 2000, media and sera for cells were from Life technologies. Oligonucleotides and most usual chemicals were from Merck-Millipore. CSA was from Clinisciences (CB0352) and AZT from the HIV reagent program (#3485).

### Recombinant proteins

Recombinant Tat (86 residues, BH10 isolate) was produced and purified from transfected *E.coli* as described (Vendeville et al., 2004) except that after induction bacteria were kept at 30°C. Recombinant human CypA was produced using a pET vector encoding a N-terminal His6 tag before a TEV protease site and CypA coding sequence (Ahmed-Belkacem et al., 2016). After growing transformed BL21 λDE3 *E. coli* cells at 37°C for 4 h, bacteria were lysed by sonication in 150 mM NaCl 150 mM, 10 mM imidazole, 1 mM DTT, 50 mM Tris, pH 8.0 supplemented with antiproteases (Complete^c^, Roche) and 1mg lysozyme / ml. The lysate was clarified by centrifugation before adding Ni-NTA-agarose (Qiagen) and incubation for 2 h at 4°C on a rotating wheel. The resin was then transferred to a column and washed with lysis buffer supplemented with 10 mM imidazole. His6-CypA was eluted with 500 mM imidazole in lysis buffer, then dialyzed overnight at 4°C against 150 mM NaCl, 50 mM Tris, pH 8.0, containing 1mM beta mercaptoethanol together with His6-TEV protease (1 mg / 50 mg of His6-CypA). Ni-NTA-agarose was then added to the dialysate to remove His6-TEV protease and any unprocessed His6-CypA. After 2 h at 4°C on a wheel and centrifugation, the supernatant containing purified CypA was collected and the purified protein that contains an extra N-terminal gly residue was stored at -80°C. Purification of the CA protein was performed as described (Hung et al., 2013). CypA-H126Q and CA-G89V pET vectors were generated using quickchange lightning (Agilent), coding sequences were entirely sequenced and proteins were purified as described above for the WT proteins.

GST-tagged proteins were purified essentially as described (Sabers et al., 1995). Briefly, E coli BL21 cells transformed with pGEX vectors containing Tat101 (Benkirane et al., 1998), CypA or CypA-H126Q (Chatterji et al., 2010) were grown at 37°C until OD=0.6. Production of GST-tagged protein was induced by adding 1 mM isopropylthiogalactoside for 4.5 h at 30°C. Bacteria were then lysed in 150 mM NaCl, 50 mM Tris pH 7.5 supplemented with antiproteases, 0.5% TX-100 and 2 mM DTT. After sonication for 1 min, bacterial extracts were centrifugated for 20 min at 18,000 x g at 4°C and the supernatant was incubated with 1.5 ml of a GSH-sepharose 4B resin (GE Healthcare 17-0756-01) for 1.5 h at 4°C on a rotating wheel. After washing with lysis buffer, beads were frozen in lysis buffer supplemented with 10% glycerol and stored as aliquots at -80°C.

### SEC experiments

CypA and CA were dialysed against gel filtration buffer (150 mM NaCl, 5 mM DTT, 50 mM citrate, pH 7.3), while lyophilized Tat was resuspended in this buffer. Proteins (18 nmoles each in 250 µl buffer) were incubated for 30 min at RT then loaded on a Superdex 200 Increase 10/300 GL column connected to an AKTA purification system (GE Healthcare). The column (bed volume 24 ml; void volume 8.8 ml) was equilibrated and eluted at 0.4 ml/min in gel filtration buffer. Fractions (0.5 ml) corresponding to chromatographic peaks were pooled before precipitation with 33% trichloroacetic acid 30 min on ice. Pellets were washed with ice-cold acetone then resuspended in reducing sample buffer before SDS/PAGE and Western blots.

### Microscale thermophoresis (MST)

Proteins were dialyzed overnight against 150 mM NaCl, 50 mM citrate, pH 7.3 the day before the experiment. CypA (WT or H126Q) were labeled with the Monolith Protein Labeling Kit RED-NHS 2^nd^ generation according to the recommendations of the manufacturer (NanoTemper Technologies). Briefly, CypA (200 µl at 10 µM) was labeled with 5 µl of labeling reagent. After 30 min the mixture was loaded on a minitrap-G25 column (GE Healthcare) and the labeled protein was eluted with 300 µl of dialysis buffer. Labeled CypA was used at 50 nM and dilutions were made in dialysis buffer containing 0.05% Tween. Protein-protein interactions were analyzed at 20°C by MST on a Monolith NT.115 (NanoTemper technologies) using standard capillaries. We made controls recommended by the manufacturer, *i.e.* absence of aggregation and interaction with capillary glass; signal homogeneity along the capillary. Measurements were repeated at least two times. K_d_ values were calculated using the NTAnalysis software (NanoTemper technologies) or GraphPad Prism.

### Surface Plasmon Resonance

The Surface Plasmon Resonance experiments were performed at 25 °C on a T200 apparatus (Cytiva) in 20 mM phosphate and 30 mM citrate, pH 7 buffer containing 150 mM NaCl and P20 surfactant 0.05% (Cytiva). CA and Tat Proteins at 50 µg/ml in 150 mM NaCl, 50 mM acetate buffer pH5 and 150 mM NaCl, citrate buffer pH 6.2 respectively were immobilized on CM5S sensor chip by amine coupling according to the manufacturer’s instructions. Around 11000 RU of Tat and 1000 RU of CA were thereby obtained. Affinities were estimated by one cycle kinetic titration at five increasing concentrations (370-6000 nM, two fold dilution series) of CypA or equimolecular CypA /CA mixing or CA injected on 50-70 RU of CypA previously bound on immobilized Tat. A two state fitting model (BIAevaluation software 4.1; Cytiva) was used for calculations. No binding of CypA was observed on CA-G89V (Figure S1).

### Cells and transfections

Cell lines were obtained from the ATCC and cultivated following their recommendations. They were checked monthly for mycoplasma contamination. Jurkat cells (ATCC clone E6-1, TIB-152) were transfected by electroporation using Amaxa kit V (program X-005), and HEK 293 T (ATCC CRL-11268) were transfected using PEImax as described (Longo et al., 2013). For the preparation of human monocytes and T-cells, human blood from healthy donors was obtained from the local blood bank (Etablissement Français du Sang, Montpellier; agreement 21PLER2019-0106). Peripheral blood mononuclear cells were isolated by separation on Ficoll-Hypaque (Eurobio). CD4+ T-cells were then purified using a CD4 Easysep negative selection kit (Stemcell). They were activated using phytohemagglutinin (1 μg/ml) for 24 h then interleukin-2 (50U/ml) for 4-6 days before nucleofection (as recommended by the manufacturer, Lonza) or infection by HIV-1.

### Infections

To produce HIV-1 virions, HEK 293 T cells were transfected with pNL4.3 using PEI max (Chopard *et al*., 2018). To prepare pseudotyped viruses, cells were cotransfected (ratio 1:5) with a pCMV-VSV-G vector and a ΔEnv pNL4.3. The latter was obtained by deletion between the NdeI (6399) and ApaLI (6610) restriction sites of pNL4.3. This deletion introduced a stop codon so that a truncated protein of 63 residues only was produced instead of the 854 residues Env protein. Vpu sequence was not altered. The ΔTat pNL4.3 clone has been described (Chopard *et al*., 2018) and ΔTat ΔEnv pNL4.3 was produced as described above.

Vpr chimera were constructed by PCR amplification of Vpr (Alfaisal et al., 2019), CypA or Tat in pCi vector (Promega) and mutations were introduced using a QuickChange lightning kit (Agilent). Coding sequences were entirely sequenced. For the complementation of the ΔTat virus, ΔTat ΔEnv pNL4.3 was cotransfected with pCMV-VSV-G (6/1 ratio) and 1/1000 of the indicated pBi-Tat vector.

For the production of viruses with Vpr chimera, HEK293T cells were cotransfected with CA-G89V ΔEnv pNL4.3, Tat-Vpr or CypA-Vpr (3/1 ratio) and pCMV-VSV-G vector (5/1 ratio). When indicated transfected cells were treated with 2 µM CSA to prevent CypA encapsidation (Franke *et al*., 1994; Thali *et al*., 1994). These viruses were termed CSA-viruses throughout this paper. The cell supernatant was harvested 48h after transfection, filtered onto 0.4 µm filters and ultracentrifugated on 20% sucrose at 125 000 x g for 2 h at 4°C. Viruses were aliquoted and stored at -80°C. Virus titers were determined by ELISA p24 (Innotest) using appropriate standards. WT CA and CA-G89V were similarly detected using this ELISA kit (Figure S3), and viral stocks were normalized using p24 ELISA. Single round infections were performed using 3-300 ng p24 of VSV-G pseudotyped viruses per million T-cell. 18h after infection, cells were washed three times in culture medium then resuspended in fresh medium that was harvested 18 h later for p24 ELISA. Negative controls included 20 µM AZT. For FACS analysis cells were stained for p24 using Kc57-RD1 or Kc57-FITC and analysed using an ACEA NovoCyte flow cytometer. This antibody similarly detected WT CA and CA-G89V (Figure S4).

### Capsid purification

Capsids were purified as described (Xu *et al*., 2020). Briefly, freshly-purified virions were resuspended in ice-cold 0.1 M MOPS (3-(N-morpholino) propane sulfonic acid), pH 7.0, and TX-100 was then added at 0.5% final concentration. After 2 min at 4°C, virions were centrifugated at 20,000 × g for 8 min at 4°C. Pellets were washed twice with 0.1M MOPS and capsids were finally resuspended in SDS/PAGE sample buffer.

### TIRF and AFM microscopy

Coverslips were first cleaned by bath sonication in 1M KOH, then rinsed and sonicated in ultrapure water. They were then treated with 10 mM NiCl_2_ for 1 h at RT. Freshly isolated viruses (WT or CA-G89V) were diluted 10-fold in PBS then applied to coverslips that were then incubated for 15 min at 37°C before fixation with 3.7 % paraformaldehyde in PBS. Following fixative neutralization with 50 mM NH_4_Cl, viruses were permeabilized with 0.05% saponin in PBS containing 1% bovine serum albumin, then labelled with rabbit anti-Tat and goat anti-p24 for 30 min, then with donkey anti-goat-alexaFluor647 and swine anti-rabbit-FITC for 30 min. Viruses were imaged within 24h with an MSNL Bruker microlever probe using an AFM microscope (Nanowizard 4, JPK Instrument) coupled with a fluorescence microscope (Axio observer, Carl Zeiss) equipped with a 63x NA1.4 objective and operated in the TIRF mode. Images were 50 µm x 50 µm (512 x 512 pixels). AFM and TIRF images were aligned using the Ec-CLEM Icy plug-in (Paul-Gilloteaux et al., 2017) and colocalization was assessed using Imaris setting a threshold of 300 nm, which is about the diffraction limit of the fluorochromes. Viruses were identified as CA^+^ TIRF spots with an AFM signal whose height >100 nm.

### qPCR and qRT-PCR

Jurkat cells were infected with VSV-G pseudotyped viruses (WT, CSA or CA-G89V; 15.3 ng p24/ million cells). Cells were pelleted for DNA extraction using DNAzol (Thermofisher) 24 h or 34 h after infection. For RNA extraction, cells were harvested 6h or 10 h after infection, washed three times with PBS containing 1% FBS, before extracting RNA using Trizol (Thermofisher). Extractions were performed following the manufacturer instructions. To monitor HIV transcription, RNA was reverse transcribed into first-strand cDNA using All-in-one RT MasterMix (Applied Biological Materials). qPCR of cDNA was then performed using P7/P8 primers (5249-5358), LightCycler 480 SYBR Green I Master mix (Roche) and normalization using GAPDH as described (Mousseau et al., 2012). To quantify reverse transcription early and late products, qPCR was performed exactly as described (Mbisa et al., 2009) using TaqMan Universal Master Mix II (Applied Biosystems). Identical amounts of DNA (∼20 ng) was used for each reaction. Standard qPCR curves ware obtained using linearized pNL4.3.

### Materials and data availability statement

Materials created for this study are freely available upon request to the authors. All data have been provided in the manuscript and supporting files, allowing research reproducibility.

## Acknowledgements

This work was funded by Sidaction (2019-AEQ-12183) to BB, FRM (Equipe FRM 20161136701) to JMM, FranceBioImaging (FBI, ANR10INSB04) and the GIS IBISA (Infrastructures en Biologie Santé et Agronomie) to PEM. We are grateful to Husam Alsarraf for help during SEC experiments, Christophe Chopard and Xavier Hanoulle for pioneer experiments, Juliette Cuminal and Melissa Ramos Mach for technical assistance, Laurent Chaloin and Jean-Marie Peloponese for expertise in structural biology and qRT-PCR, respectively.

## Author contributions

MS, CO, LM, IC performed *in cellulo* proteins interaction studies. MS, CO performed SEC analysis supervised by MB. MS, JFG and MB performed thermophoresis analysis. MP and CH analyzed interactions using Biacore. VT, CO and LM analyzed purified virions and capsids. MS, LC, CG and PEM performed TIRF/AFM experiments. CM prepared all HIV mutants. MS, CO, LM and BB performed infectivity studies. JMM provided funding and scientific input. BB supervised the study and wrote the manuscript with inputs from all the authors.

The authors declare that they have no conflict of interest.

---

**Figure S1.**
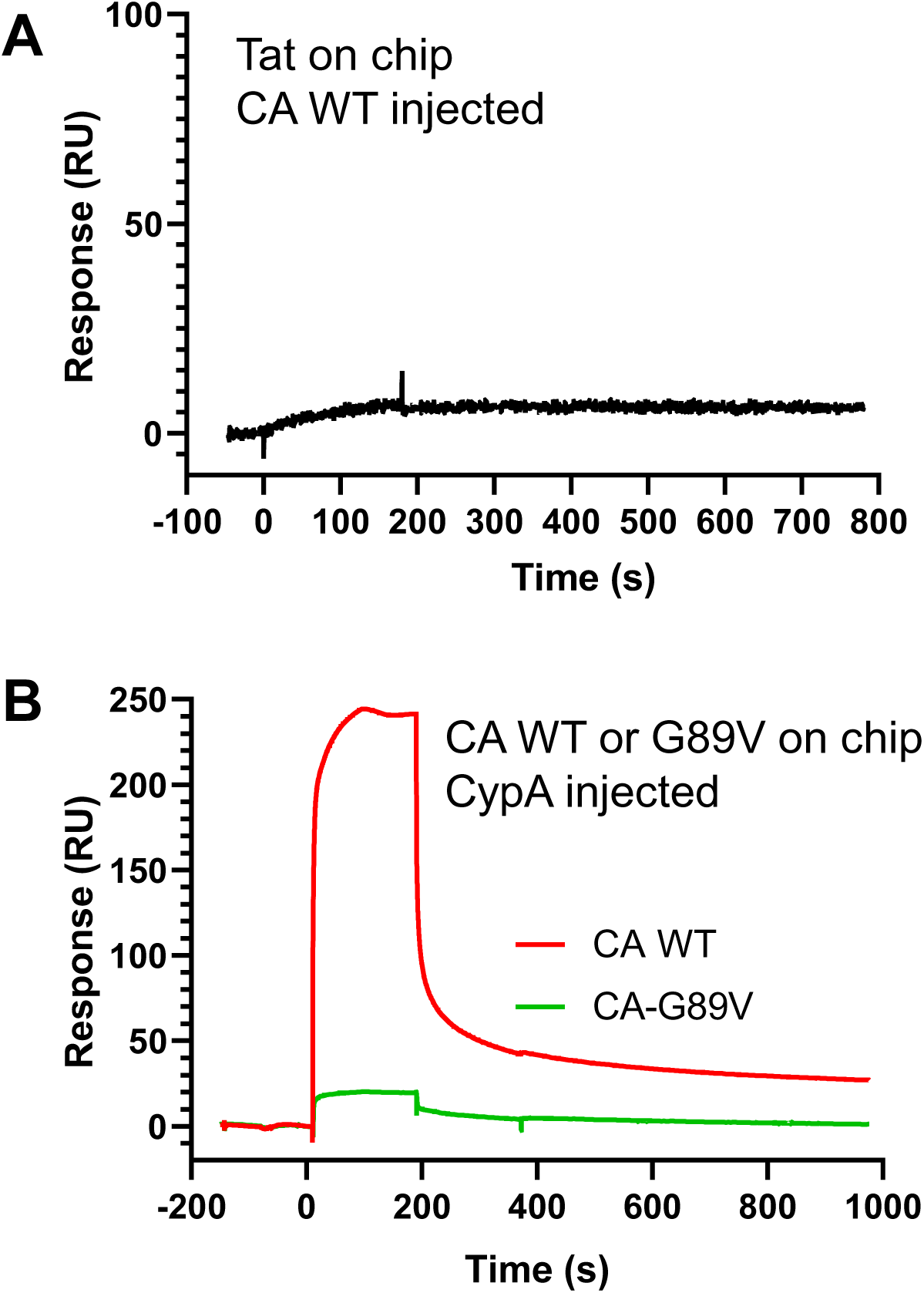
SPR analysis. A, Tat does not directly interact with CA. Tat was immobilized on a chip before applying 2 µM CA WT. B, CypA binds to CA WT but not CA-G89V. Capsid proteins were immobilized on a chip before applying 15 µM CypA. **Figure supplement-source data.** Datasheets for graphs.

**Figure S2.**
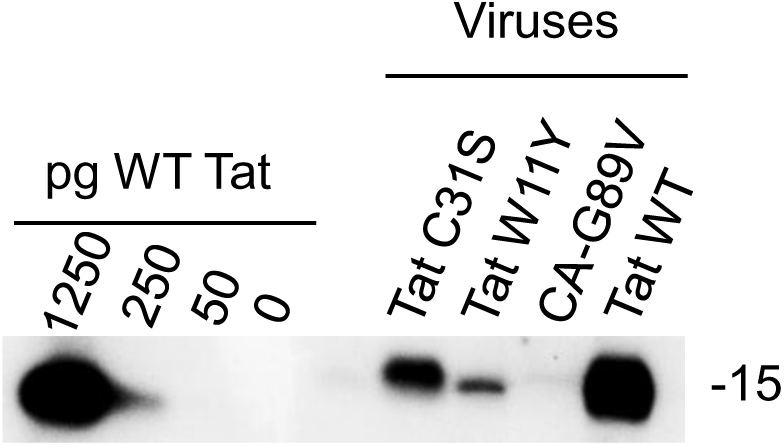
Semiquantitative blot of purified viruses. Proteins from WT or mutant viruses and recombinant Tat standards (101 residues) were separated on 16 % Tricine gels before anti-Tat western blot. **Figure supplement-source data. R**aw immunoblot.

**Figure S3.**
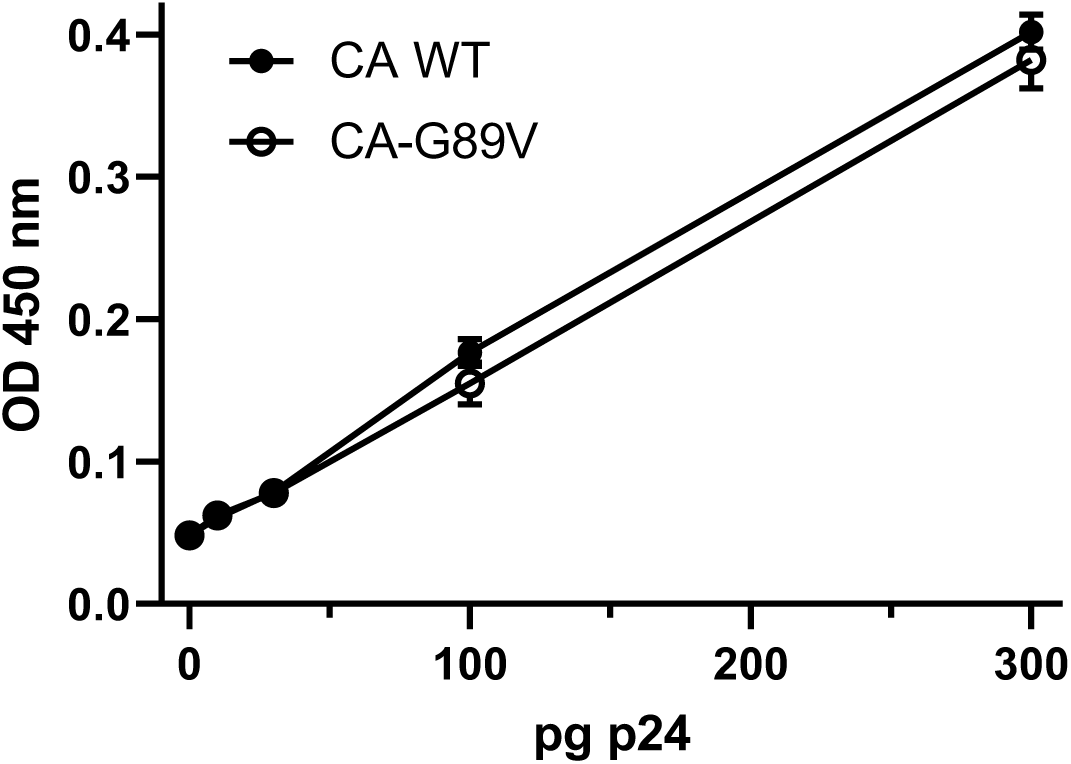
WT and G89V capsids are similarly detected using p24 ELISA. HIV-1 capsid proteins were produced and purified from *E. coli* as described in methods and ELISA was performed as recommended by the manufacturer. Means ± SEM (n=2). Most error bars are within the symbol size. **Figure supplement-source data.** Datasheet for the graph.

**Figure S4.**
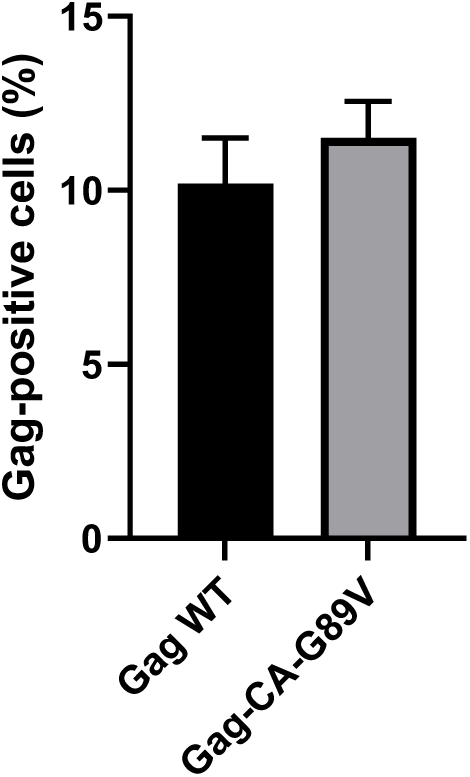
Gag-WT and Gag-CA-G89V capsids are similarly detected by FACS analysis. Jurkat cells were transfected with Gag-WT or Gag-CA-G89V as indicated. Cells were washed then fixed and permeabilized before staining with Kc57-FITC and analysis by FACS. Means ± SEM (n=2). **Figure supplement-source data.** Datasheet for the graph.

Sources of antibodies
| Antigen | Ab species | Label | source | reference |
| --- | --- | --- | --- | --- |
| Tat | Mouse monoclonal |  | SantaCruz Biotechnology | sc65912 |
| Tat | Rabbit |  | E. Loret | home made |
| Tat | Mouse monoclonal |  | E. Loret | 7G12 |
| CypA | Rabbit |  | SantaCruz Biotechnology | sc133494 |
| CypA | Goat |  | R/D Systems | AF3589 |
| HIV p24 (CA) | Goat |  | BioRad | 4999-9007 |
| HIV p24 (CA) | Mouse monoclonal |  | AIDS Research and Reference Reagent Program | 183-H12-5C |
| HIV p24 (CA) | Mouse monoclonal | RD1 | Beckman Coulter | Kc57-RD1 |
| FKBP12 | Rabbit |  | Pierce | PA1-026A |
| Mouse IgG | Goat | HRP | Jackson ImmunoResearch | 115-035-146 |
| Rabbit IgG | Goat | HRP | Jackson ImmunoResearch | 111-035-144 |
| Rabbit IgG | Swine | FITC | Nordic | F0205 |
| Goat IgG | Rabbit | HRP | Sigma | A-5420 |
| Goat IgG | Donkey | AlexaFluor647 | Invitrogen | A21447 |

Antibodies are polyclonal except when otherwise indicated

## References

Ahmed-Belkacem, A., Colliandre, L., Ahnou, N., Nevers, Q., Gelin, M., Bessin, Y., Brillet, R., Cala, O., Douguet, D., Bourguet, W., et al. (2016). Fragment-based discovery of a new family of non-peptidic small-molecule cyclophilin inhibitors with potent antiviral activities. Nature communications 7, 12777.

Alfaisal, J., Machado, A., Galais, M., Robert-Hebmann, V., Arnaune-Pelloquin, L., Espert, L., and Biard-Piechaczyk, M. (2019). HIV-1 Vpr inhibits autophagy during the early steps of infection of CD4 T cells. Biol Cell 111, 308–318. 10.1111/boc.201900071.

Apolloni, A., Hooker, C.W., Mak, J., and Harrich, D. (2003). Human immunodeficiency virus type 1 protease regulation of tat activity is essential for efficient reverse transcription and replication. J Virol 77, 9912–9921. 10.1128/jvi.77.18.9912-9921.2003.

Bayer, P., Kraft, M., Ejchart, A., Westendorp, M., Frank, R., and Rosch, P. (1995). Structural studies of HIV-1 Tat protein. J Mol Biol 247, 529–535.

Benkirane, M., Chun, R.F., Xiao, H., Ogryzko, V.V., Howard, B.H., Nakatani, Y., and Jeang, K.T. (1998). Activation of integrated provirus requires histone acetyltransferase. p300 and P/CAF are coactivators for HIV-1 Tat. J Biol Chem 273, 24898–24905.

Boudier, C., Humbert, N., Chaminade, F., Chen, Y., de Rocquigny, H., Godet, J., Mauffret, O., Fosse, P., and Mely, Y. (2014). Dynamic interactions of the HIV-1 Tat with nucleic acids are critical for Tat activity in reverse transcription. Nucleic Acids Res 42, 1065–1078.

Braaten, D., Franke, E.K., and Luban, J. (1996). Cyclophilin A is required for an early step in the life cycle of human immunodeficiency virus type 1 before the initiation of reverse transcription. Journal of virology 70, 3551–3560.

Briggs, C.J., Ott, D.E., Coren, L.V., Oroszlan, S., and Tozser, J. (1999). Comparison of the effect of FK506 and cyclosporin A on virus production in H9 cells chronically and newly infected by HIV-1. Arch Virol 144, 2151–2160. 10.1007/s007050050629.

Butler, S.L., Hansen, M.S., and Bushman, F.D. (2001). A quantitative assay for HIV DNA integration in vivo. Nature medicine 7, 631–634.

Chapman, E.H., Kurec, A.S., and Davey, F.R. (1981). Cell volumes of normal and malignant mononuclear cells. J Clin Pathol 34, 1083–1090. 10.1136/jcp.34.10.1083.

Chatterji, U., Lim, P., Bobardt, M.D., Wieland, S., Cordek, D.G., Vuagniaux, G., Chisari, F., Cameron, C.E., Targett-Adams, P., Parkinson, T., and Gallay, P.A. (2010). HCV resistance to cyclosporin A does not correlate with a resistance of the NS5A-cyclophilin A interaction to cyclophilin inhibitors. J Hepatol 53, 50–56. 10.1016/j.jhep.2010.01.041.

Chertova, E., Chertov, O., Coren, L.V., Roser, J.D., Trubey, C.M., Bess, J.W., Jr., Sowder, R.C., 2nd, Barsov, E., Hood, B.L., Fisher, R.J., et al. (2006). Proteomic and biochemical analysis of purified human immunodeficiency virus type 1 produced from infected monocyte-derived macrophages. J Virol 80, 9039–9052.

Chopard, C., Tong, P.B.V., Toth, P., Schatz, M., Yezid, H., Debaisieux, S., Mettling, C., Gross, A., Pugniere, M., Tu, A., et al. (2018). Cyclophilin A enables specific HIV-1 Tat palmitoylation and accumulation in uninfected cells. Nature communications 9, 2251.

Dahmane S, Doucet C, Le Gall A, Chamontin C, Dosset P, Murcy F, Fernandez L, Salas D, Rubinstein E, Mougel M, et al. (2019). Nanoscale organization of tetraspanins during HIV-1 budding by correlative dSTORM/AFM. Nanoscale 11, 6036–6044. 10.1039/c8nr07269h.

De Iaco, A., and Luban, J. (2014). Cyclophilin A promotes HIV-1 reverse transcription but its effect on transduction correlates best with its effect on nuclear entry of viral cDNA. Retrovirology 11, 11.

Ensoli, B., Barillari, G., Salahuddin, S.Z., Gallo, R.C., and Wong-Staal, F. (1990). Tat protein of HIV-1 stimulates growth of cells derived from Kaposi’s sarcoma lesions of AIDS patients. Nature 345, 84–86.

Feinberg, M.B., Baltimore, D., and Frankel, A.D. (1991). The role of Tat in the human immunodeficiency virus life cycle indicates a primary effect on transcriptional elongation. Proc Natl Acad Sci U S A 88, 4045–4049. 10.1073/pnas.88.9.4045.

Fernandez, J., Machado, A.K., Lyonnais, S., Chamontin, C., Gartner, K., Leger, T., Henriquet, C., Garcia, C., Portilho, D.M., Pugniere, M., et al. (2019). Transportin-1 binds to the HIV-1 capsid via a nuclear localization signal and triggers uncoating. Nat Microbiol 4, 1840–1850. 10.1038/s41564-019-0575-6.

Foster, T.L., Gallay, P., Stonehouse, N.J., and Harris, M. (2011). Cyclophilin A interacts with domain II of hepatitis C virus NS5A and stimulates RNA binding in an isomerase-dependent manner. J Virol 85, 7460–7464. 10.1128/JVI.00393-11.

Francis, A.C., and Melikyan, G.B. (2018). Single HIV-1 Imaging Reveals Progression of Infection through CA-Dependent Steps of Docking at the Nuclear Pore, Uncoating, and Nuclear Transport. Cell Host Microbe 23, 536–548 e536. 10.1016/j.chom.2018.03.009.

Franke, E.K., Yuan, H.E., and Luban, J. (1994). Specific incorporation of cyclophilin A into HIV-1 virions. Nature 372, 359–362.

Freed, E.O. (2015). HIV-1 assembly, release and maturation. Nature reviews Microbiology 13, 484–496.

Harrich, D., Ulich, C., Garcia-Martinez, L.F., and Gaynor, R.B. (1997). Tat is required for efficient HIV-1 reverse transcription. Embo J 16, 1224–1235.

Howard, B.R., Vajdos, F.F., Li, S., Sundquist, W.I., and Hill, C.P. (2003). Structural insights into the catalytic mechanism of cyclophilin A. Nat Struct Biol 10, 475–481.

Hung, M., Niedziela-Majka, A., Jin, D., Wong, M., Leavitt, S., Brendza, K.M., Liu, X., and Sakowicz, R. (2013). Large-scale functional purification of recombinant HIV-1 capsid. PLoS One 8, e58035. 10.1371/journal.pone.0058035.

Jeang, K.T., Xiao, H., and Rich, E.A. (1999). Multifaceted activities of the HIV-1 transactivator of transcription, Tat. J Biol Chem 274, 28837–28840.

Karn, J., and Stoltzfus, C.M. (2012). Transcriptional and posttranscriptional regulation of HIV-1 gene expression. Cold Spring Harb Perspect Med 2, a006916. 10.1101/cshperspect.a006916.

Kim, K., Dauphin, A., Komurlu, S., McCauley, S.M., Yurkovetskiy, L., Carbone, C., Diehl, W.E., Strambio-De-Castillia, C., Campbell, E.M., and Luban, J. (2019). Cyclophilin A protects HIV-1 from restriction by human TRIM5alpha. Nat Microbiol 4, 2044–2051. 10.1038/s41564-019-0592-5.

Lahaye, X., Satoh, T., Gentili, M., Cerboni, S., Silvin, A., Conrad, C., Ahmed-Belkacem, A., Rodriguez, Elisa C., Guichou, J.-F., Bosquet, N., et al. (2016). Nuclear Envelope Protein SUN2 Promotes Cyclophilin-A-Dependent Steps of HIV Replication. Cell Reports 15, 879–892. 10.1016/j.celrep.2016.03.074.

Lalonde, M.S., Lobritz, M.A., Ratcliff, A., Chamanian, M., Athanassiou, Z., Tyagi, M., Wong, J., Robinson, J.A., Karn, J., Varani, G., and Arts, E.J. (2011). Inhibition of both HIV-1 reverse transcription and gene expression by a cyclic peptide that binds the Tat-transactivating response element (TAR) RNA. PLoS pathogens 7, e1002038.

Longo, P.A., Kavran, J.M., Kim, M.-S., and Leahy, D.J. (2013). Transient mammalian cell transfection with polyethylenimine (PEI). Methods in enzymology 529, 227–240.

Luban, J., Bossolt, K.L., Franke, E.K., Kalpana, G.V., and Goff, S.P. (1993). Human immunodeficiency virus type 1 Gag protein binds to cyclophilins A and B. Cell 73, 1067–1078.

Marchio, S., Alfano, M., Primo, L., Gramaglia, D., Butini, L., Gennero, L., De Vivo, E., Arap, W., Giacca, M., Pasqualini, R., and Bussolino, F. (2005). Cell surface-associated Tat modulates HIV-1 infection and spreading through a specific interaction with gp120 viral envelope protein. Blood 105, 2802–2811.

Mbisa, J.L., Delviks-Frankenberry, K.A., Thomas, J.A., Gorelick, R.J., and Pathak, V.K. (2009). Real-time PCR analysis of HIV-1 replication post-entry events. Methods Mol Biol 485, 55–72. 10.1007/978-1-59745-170-3_5.

McMahon, M.A., Shen, L., and Siliciano, R.F. (2009). New approaches for quantitating the inhibition of HIV-1 replication by antiviral drugs in vitro and in vivo. Curr Opin Infect Dis 22, 574–582. 10.1097/QCO.0b013e328332c54d.

Mediouni, S., Watkins, J.D., Pierres, M., Bole, A., Loret, E.P., and Baillat, G. (2012). A monoclonal antibody directed against a conformational epitope of the HIV-1 trans-activator (Tat) protein neutralizes cross-clade. The Journal of biological chemistry 287, 11942–11950.

Mousseau, G., Clementz, M.A., Bakeman, W.N., Nagarsheth, N., Cameron, M., Shi, J., Baran, P., Fromentin, R., Chomont, N., and Valente, S.T. (2012). An analog of the natural steroidal alkaloid cortistatin A potently suppresses Tat-dependent HIV transcription. Cell host & microbe 12, 97–108.

Ott, M., Geyer, M., and Zhou, Q. (2011). The control of HIV transcription: keeping RNA polymerase II on track. Cell Host Microbe 10, 426–435.

Paul-Gilloteaux, P., Heiligenstein, X., Belle, M., Domart, M.C., Larijani, B., Collinson, L., Raposo, G., and Salamero, J. (2017). eC-CLEM: flexible multidimensional registration software for correlative microscopies. Nat Methods 14, 102–103. 10.1038/nmeth.4170.

Piotukh, K., Gu, W., Kofler, M., Labudde, D., Helms, V., and Freund, C. (2005). Cyclophilin A binds to linear peptide motifs containing a consensus that is present in many human proteins. The Journal of biological chemistry 280, 23668–23674.

Rayne, F., Debaisieux, S., Yezid, H., Lin, Y.L., Mettling, C., Konate, K., Chazal, N., Arold, S.T., Pugniere, M., Sanchez, F., et al. (2010). Phosphatidylinositol-(4,5)-bisphosphate enables efficient secretion of HIV-1 Tat by infected T-cells. EMBO J 29, 1348–1362.

Sabers, C.J., Martin, M.M., Brunn, G.J., Williams, J.M., Dumont, F.J., Wiederrecht, G., and Abraham, R.T. (1995). Isolation of a protein target of the FKBP12-rapamycin complex in mammalian cells. J Biol Chem 270, 815–822. 10.1074/jbc.270.2.815.

Schaller, T., Ocwieja, K.E., Rasaiyaah, J., Price, A.J., Brady, T.L., Roth, S.L., Hue, S., Fletcher, A.J., Lee, K., KewalRamani, V.N., et al. (2011). HIV-1 capsid-cyclophilin interactions determine nuclear import pathway, integration targeting and replication efficiency. PLoS pathogens 7, e1002439.

Selyutina, A., Persaud, M., Simons, L.M., Bulnes-Ramos, A., Buffone, C., Martinez-Lopez, A., Scoca, V., Di Nunzio, F., Hiatt, J., Marson, A., et al. (2020). Cyclophilin A Prevents HIV-1 Restriction in Lymphocytes by Blocking Human TRIM5alpha Binding to the Viral Core. Cell Rep 30, 3766–3777 e3766. 10.1016/j.celrep.2020.02.100.

Sokolskaja, E., Berthoux, L., and Luban, J. (2006). Cyclophilin A and TRIM5alpha independently regulate human immunodeficiency virus type 1 infectivity in human cells. J Virol 80, 2855–2862. 10.1128/JVI.80.6.2855-2862.2006.

Sokolskaja, E., Sayah, D.M., and Luban, J. (2004). Target cell cyclophilin A modulates human immunodeficiency virus type 1 infectivity. Journal of virology 78, 12800–12808.

Sundquist, W.I., and Krausslich, H.-G. (2012). HIV-1 assembly, budding, and maturation. Cold Spring Harb Perspect Med 2, a006924.

Swanson, C.M., and Malim, M.H. (2008). SnapShot: HIV-1 proteins. Cell 133, 742, 742.e741.

Thali, M., Bukovsky, A., Kondo, E., Rosenwirth, B., Walsh, C.T., Sodroski, J., and Gottlinger, H.G. (1994). Functional association of cyclophilin A with HIV-1 virions. Nature 372, 363–365. 10.1038/372363a0.

Towers, G.J., Hatziioannou, T., Cowan, S., Goff, S.P., Luban, J., and Bieniasz, P.D. (2003). Cyclophilin A modulates the sensitivity of HIV-1 to host restriction factors. Nat Med 9, 1138–1143.

Vendeville, A., Rayne, F., Bonhoure, A., Bettache, N., Montcourrier, P., and Beaumelle, B. (2004). HIV-1 Tat enters T cells using coated pits before translocating from acidified endosomes and eliciting biological responses. Mol Biol Cell 15, 2347–2360.

Verhoef, K., Koper, M., and Berkhout, B. (1997). Determination of the minimal amount of Tat activity required for human immunodeficiency virus type 1 replication. Virology 237, 228–236. 10.1006/viro.1997.8786.

Xu, C., Fischer, D.K., Rankovic, S., Li, W., Dick, R.A., Runge, B., Zadorozhnyi, R., Ahn, J., Aiken, C., Polenova, T., et al. (2020). Permeability of the HIV-1 capsid to metabolites modulates viral DNA synthesis. PLoS Biol 18, e3001015. 10.1371/journal.pbio.3001015.

Yao, X.J., Kobinger, G., Dandache, S., Rougeau, N., and Cohen, E. (1999). HIV-1 Vpr-chloramphenicol acetyltransferase fusion proteins: sequence requirement for virion incorporation and analysis of antiviral effect. Gene Ther 6, 1590–1599.

Yoo, S., Myszka, D.G., Yeh, C., McMurray, M., Hill, C.P., and Sundquist, W.I. (1997). Molecular recognition in the HIV-1 capsid/cyclophilin A complex. J Mol Biol 269, 780–795.

Zhang, J., Tamilarasu, N., Hwang, S., Garber, M.E., Huq, I., Jones, K.A., and Rana, T.M. (2000). HIV-1 TAR RNA enhances the interaction between Tat and cyclin T1. J Biol Chem 275, 34314–34319.

Zila, V., Margiotta, E., Turonova, B., Muller, T.G., Zimmerli, C.E., Mattei, S., Allegretti, M., Borner, K., Rada, J., Muller, B., et al. (2021). Cone-shaped HIV-1 capsids are transported through intact nuclear pores. Cell 184, 1032–1046 e1018. 10.1016/j.cell.2021.01.025.

